# Latitudinal scaling of aggregation with abundance and its consequences for coexistence in species rich forests

**DOI:** 10.1101/2023.05.18.541254

**Authors:** Thorsten Wiegand, Xugao Wang, Samuel M. Fischer, Nathan J. B. Kraft, Norman A. Bourg, Warren Y. Brockelman, Guanghong Cao, Min Cao, Wirong Chanthorn, Chengjin Chu, Stuart Davies, Sisira Ediriweera, C. V. S. Gunatilleke, I. A. U. N. Gunatilleke, Zhanqing Hao, Robert Howe, Mingxi Jiang, Guangze Jin, W. John Kress, Buhang Li, Juyu Lian, Luxiang Lin, Feng Liu, Keping Ma, William McShea, Xiangcheng Mi, Jonathan A. Myers, Anuttara Nathalang, David A. Orwig, Guochun Shen, Sheng-Hsin Su, I-Fang Sun, Xihua Wang, Amy Wolf, Enrong Yan, Wanhui Ye, Yan Zhu, Andreas Huth

## Abstract

The search for simple principles underlying the complex spatial structure and dynamics of plant communities is a long-standing challenge in ecology^1-6^. In particular, the relationship between the spatial distribution of plants and species coexistence is challenging to resolve in species-rich communities^7-9^. Analysing the spatial patterns of tree species in 21 large forest plots, we find that rare species tend to be more spatially aggregated than common species, and a latitudinal gradient in the strength of this negative correlations that increases from tropical to temperate forests. Our analysis suggests that latitudinal gradients in animal seed dispersal^10^ and mycorrhizal associations^11,12,13^ may jointly generate this intriguing pattern. To assess the consequences of negative aggregation-abundance correlations for species coexistence, we present here a framework to incorporate the observed spatial patterns into population models^8^ along with an analytical solution for the local extinction risk^14^ of species invading from low abundances in dependence of spatial structure, demographic parameters, and immigration. For example, the stabilizing effect of the observed spatial patterns reduced the local extinction risk of species when rare almost by a factor of two. Our approach opens up new avenues for integrating observed spatial patterns into mathematical theory, and our findings demonstrate that spatial patterns, such as species aggregation and segregation, can contribute substantially to coexistence in species-rich communities. This underscores the need to understand the interactions between multiple ecological processes and spatial patterns in greater detail.

## Main Text

Tropical forests have been investigated by ecologists for decades, but their high species richness still remains a key challenge for ecological theory^1-6,15^. Although a large number of studies have been devoted to this issue, mechanistic connections among key features of forests and species coexistence are not understood^7-9^. One key feature is the widespread spatial aggregation of tree species, which has been used for long to infer mechanisms contributing to coexistence in tropical forests^16-21^ because aggregation is related to ecological processes such as negative conspecific density dependence^16,17,21,22^, dispersal limitation^2,23^, distance-decay^20^, mycorrhizal associations^11,12,13^, and habitat filtering^24,25^. Conspecific spatial aggregation Ω is usually defined as the average density *D* of conspecific trees in neighbourhoods around each individual tree, divided by the mean tree density *λ* of the species in the plot^19,25^. Hence, Ω describes the extent to which trees of the same species tend to occur in spatial clusters. Theoretical and observational studies suggest that conspecific aggregation should be related to species abundance, with rarer species showing higher levels of aggregation^8,18,19,20,26,27,28^, whereas other studies find only weak relationships between aggregation and species abundance^29^ (Supplementary Text).

From a theoretical perspective, different aggregation-abundance relationships are possible. At one extreme, when clusters form mostly near adults (e.g., due to short-distance dispersal^7,8,26^ and/or mycorrhizal associations^11,12,13^), the density *D* of conspecific trees in the neighbourhood of individual trees is similar for rare and common species, but rarer species have fewer clusters (Fig. 1a versus 1b). Thus, aggregation (i.e., Ω=*D/λ*^19,25^) increases if abundance and therefore mean tree density *λ* decrease. At the other extreme, when local clusters are created away from conspecific adults^8^ (e.g., due to clumped animal seed dispersal, canopy gaps or edaphic habitat conditions^24,30,31,32,33^), fewer seeds will reach the cluster locations when the species is rarer. This makes the neighbourhood density *D* proportional to abundance and thus aggregation Ω independent of abundance (Fig. 1a versus 1c).

**Fig. 1.**
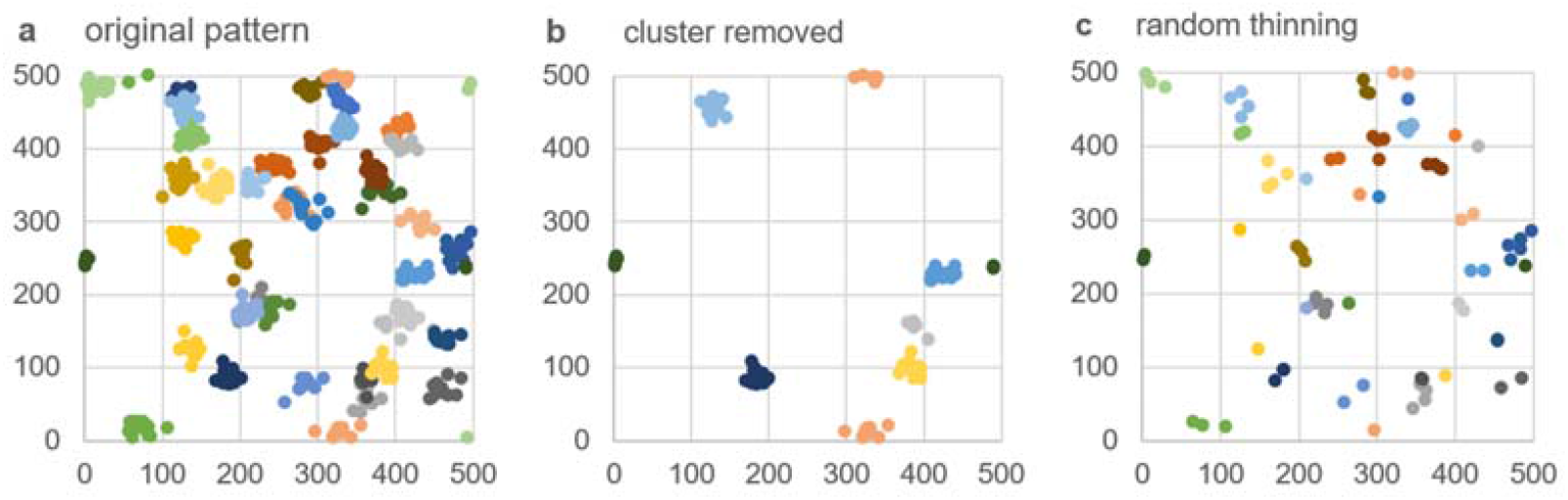
Different responses of conspecific spatial aggregation to changes in abundance. **a**, clustered pattern of an abundant species with *N* = 500 individuals with mean neighbourhood density *D* = 0.012, mean tree density *λ*= 0.002, and aggregation Ω = *D*/*λ* = 5.8 (colours represent different clusters of individuals of the same species), **b**, entire clusters of the pattern in a) were removed (*N* = 100) to maintain the neighbourhood density *D*, but since *λ* was reduced by factor 1/5, aggregation increased 5 times (Ω = 26.9). **c**, 400 individuals of the pattern in a) were randomly removed to approximately maintain aggregation Ω = 4.8 (i.e., both, *D* and *λ* were reduced by the same factor 1/5).

Given the links between conspecific aggregation, negative density dependence and coexistence^7,8,9,22,26^, the aggregation-abundance relationship may be related to the latitudinal diversity gradient^34^. Here, we conduct a comprehensive analysis of how spatial neighbourhood patterns of trees derived from large forest inventories^6^ and the relationship of aggregation and abundance change with latitude. We propose underlying mechanisms, and integrate our results into mathematical theory to investigate how the aggregation-abundance relationship may affect species co-existence (Box 1).

While aggregation can be defined in a variety of ways^18,19,25,29,35^, we use an up-scaling approach^8^ to derive biologically motivated measures of spatial patterns (such as aggregation) from established approaches to model the effect of neighbours on the performance of individual plants^36,37^ (Box 1). We exemplify our new theory for neighbourhood crowding competition, where tree survival is reduced in areas of high tree neighbourhood density via competition for space, light or nutrients^38^, or through natural enemies^16,17^.

### Box 1.

**Crowding indices, aggregation, segregation, and their link with macroscale dynamics**

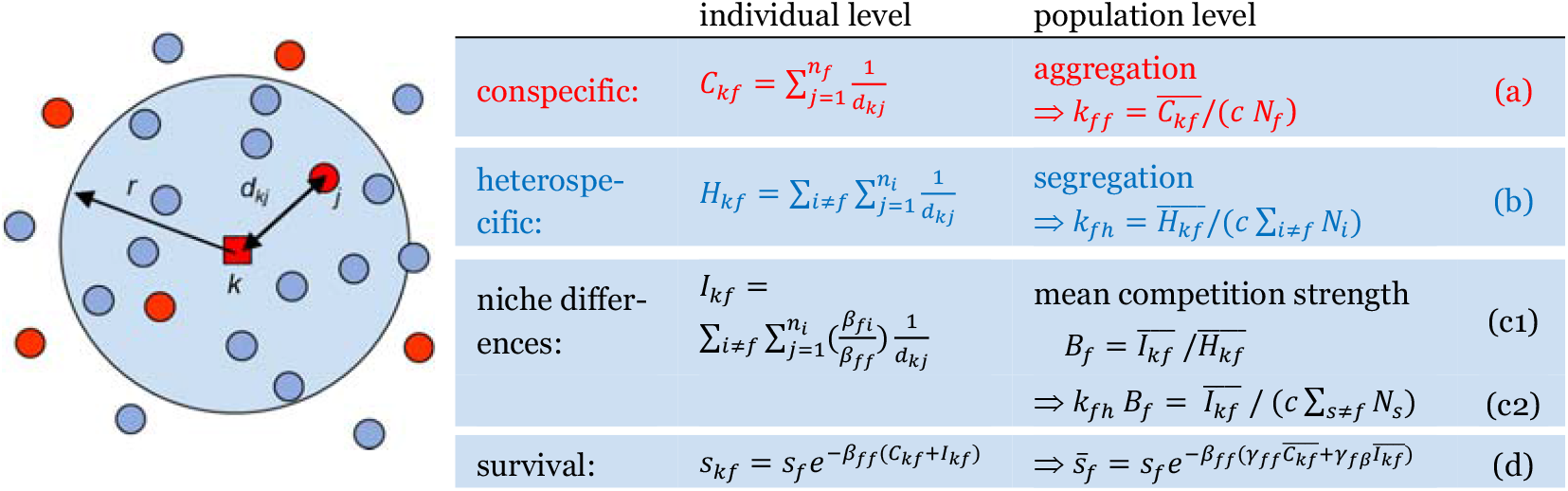

Following earlier work on **neighbourhood crowding indices** (*NCI*’s^36,37^), we assume that the survival of an individual (red square) depends on the proximity of conspecific (red) and heterospecific (blue) neighbours within its neighbourhood of radius *r* (blue shaded area). The crowding index *C*_*kf*_ counts all conspecific individuals *j* with distances *d*_*kj*_ < *r* of the focal individual *k*, but weights them by 1/*d*_*kj*_ (eq. a), assuming that distant neighbours compete less. The crowding index *H*_*kf*_ does the same with all heterospecifics (eq. b), and the crowding index *I*_*kf*_ weights neighbours of species *i* additionally by their relative competition strength *β*_*fi*_/*β*_*ff*_ (eq. c1), where ***β***_***fi***_ **is the individual-level interaction coefficient**, which measures the negative impact of one neighbour of species *i* on survival of individuals of the focal species *f*. Total crowding *C*_*kf*_ + *I*_*kf*_ then governs the survival of the focal individual^36^ (eq. d).

We find that the **population averages** 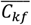 **and** 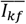 of the crowding indices determine the **average survival rate** 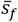 of species *f* (eq. d; eq. 5 in methods). To incorporate the average survival rate into a model of community dynamics (eq. 7 in methods), we decompose 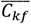 and 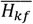 into species abundance *N*_*f*_ and measures of spatial patterns (eqs. a, b, c2; eq. 6 in methods). **Conspecific aggregation** *k*_*ff*_ describes the extent to which trees of the same species tend to occur in spatial clusters and **heterospecific segregation** *k*_*fh*_ describes the extent to which they are separated from heterospecifics. We define our measures *k*_*ff*_ and *k*_*fh*_ by dividing the average crowding indices 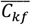 and 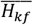 by their expectation in the absence of spatial patterns^19,25^ (eqs. a, b), where *c* is a scaling factor (*c* = 2 π *r*/*A*, see methods) and *A* is the area of the plot.

The emerging quantity *B*_*f*_ (eq. c1) is the **average competition strength of one heterospecific neighbour** relative to that of one conspecific. We compare two scenarios for the individual-level interaction coefficient *β*_*fi*_, one where con- and heterospecifics compete equally (*β*_*fi*_*/β*_*ff*_ = 1), and one with phylogenetic similarity as proxy for *β*_*fi*_/*β*_*ff*_, because it is difficult to estimate *β*_*fi*_/*β*_*ff*_ for species rich forests in practise ^37^.

Using data of 720 focal species in 21 large temperate, subtropical and tropical forest plots with sizes of 20 – 50 ha from a global network of forest research plots (CTFS-ForestGEO^6^) (Supplementary Table 1), we find that rare species tend to be more aggregated than common species (Fig. 2, Extended Data Fig. 1). Strikingly, when fitting the data of each forest plot to a power law^20,27,28^, we find that the exponent follows a strong latitudinal gradient (Fig. 2c, Extended Data Fig. 1). Tropical forests show only weak negative correlations between aggregation and abundance (Fig. 1a), but in temperate forests aggregation generally increases strongly with decreasing abundance (Fig. 2b). In contrast to conspecifics, heterospecific segregation was not correlated with abundance (except for some weaker correlations at temperate forests; Extended Data Fig. 2c, d).

**Fig. 2.**
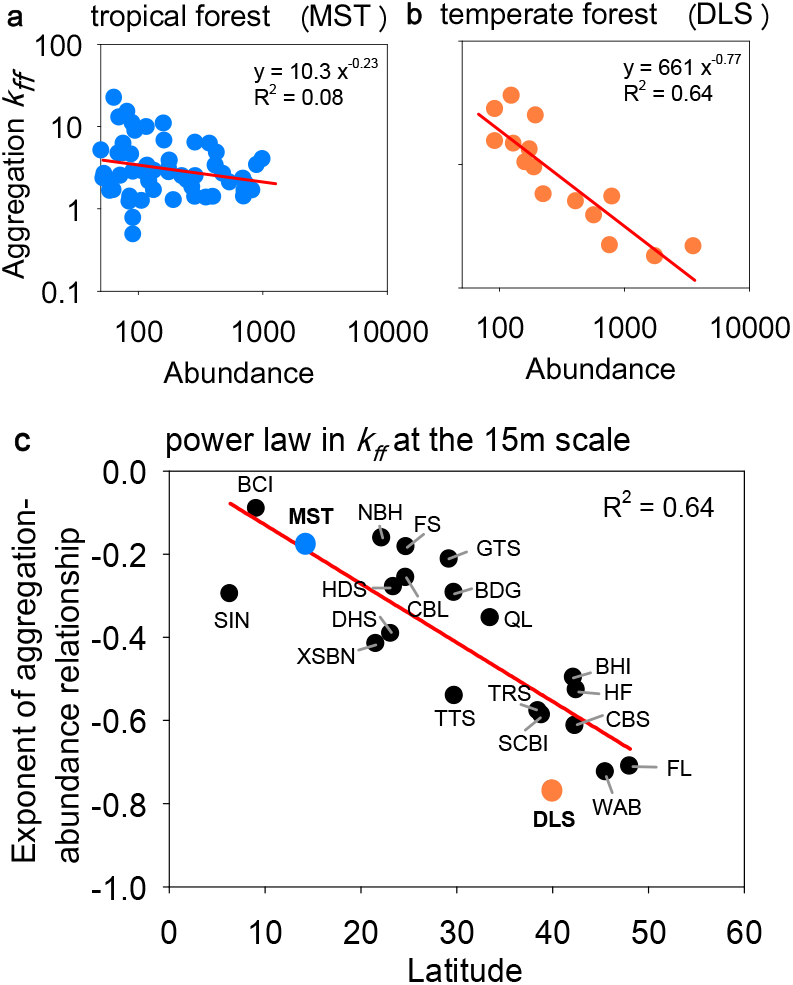
Latitudinal variation in the scaling of conspecific aggregation with abundance. **a**, aggregation values for the species in a tropical forest (Mo Singto plot; MST) plotted over their respective abundances (dots) and fitted relationship (line), **b**, same as a), but for a temperate forest (Donglingshan plot; DLS), **c**, latitudinal gradient in the exponent *b*_*f*_ of the aggregation-abundance relationship for the 21 forest plots. Aggregation is defined in Box 1 (eq. a) and is estimated for neighbourhoods of 15 m. We use in our analyses 720 species with at least 50 large trees^19^ (diameter at breast height (dbh) ≥ 10 cm. For plot characteristics and acronyms see Supplementary Table 1.

For tropical forests (i.e., low latitudes), our results indicate that mean conspecific neighbourhood tree densities 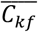 increase almost linearly with species abundance (as leaving aggregation *k*_*ff*_ constant leads to 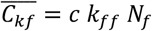; eq. a in Box 1; Fig. 1b). Thus, individuals of rare species will seldom experience the effects of intraspecific competition in this scenario. However, for temperate forests (i.e., higher latitudes), the exponent *b*_*f*_ approaches values of −1, and local conspecific neighbourhood densities are almost independent of abundance (from 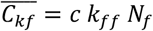 and *k*_*ff*_ = *a*_*f*_/*N*_*f*_ follows 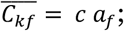 eq. a in Box 1; Fig. 1c). Thus, individuals of rare and common species experience similar degrees of conspecific competition in these forests. This counteracts a rare species advantage and as a consequence, existing theory based on invasion analysis can break down^9^. Thus, we find that spatial aggregation of trees is related to species coexistence in important ways, and we need new theories to determine whether and under which circumstances rare species are likely to increase in abundance^9^.

The latitudinal gradient in the relationship between conspecific aggregation and abundance (Fig. 1c) and the absence of such a relationship for heterospecifics (Extended Data Fig. 2c) suggest that simple principles may drive the complex spatial structure and dynamics of plant communities across latitudinal gradients. We also found similar latitudinal gradients in the proportion of species showing mostly animal seed dispersal^10^, as well as the proportion of species with an arbuscular mycorrhizal (AM) association^13^. Temperate forests are dominated by EM tree species and tropical forests by AM tree species (Extended Data Fig. 3b, c).

For the combination of the two traits (i.e., zoochory and AM association), we find an even stronger latitudinal gradient (Fig. 3a), suggesting an explanation of the observed latitudinal gradient in the aggregation-abundance relationship (Fig. 3b). Seed dispersal close to conspecific adults should be advantageous in temperate forests where species-specific EM fungi facilitate conspecific recruitment via root protection^13,39^ and counteract negative enemy-driven effects^11,12,13^. In contrast, seed dispersal farther away from conspecific adults should be more advantageous for AM trees, given that an AM association offers less protection from enemies that accumulate near conspecifics than does an EM association^12,39^. In summary, the key mechanism that leads to the different responses of aggregation to abundance is the way how local clusters emerge with respect to conspecific adults: in tropical forests, mechanisms such as animal seed dispersal lead to emergence of clusters away from adults (Fig. 1a, c), but in temperate forests clusters form close to adults (Fig. 1a, b)^8^.

**Fig. 3.**
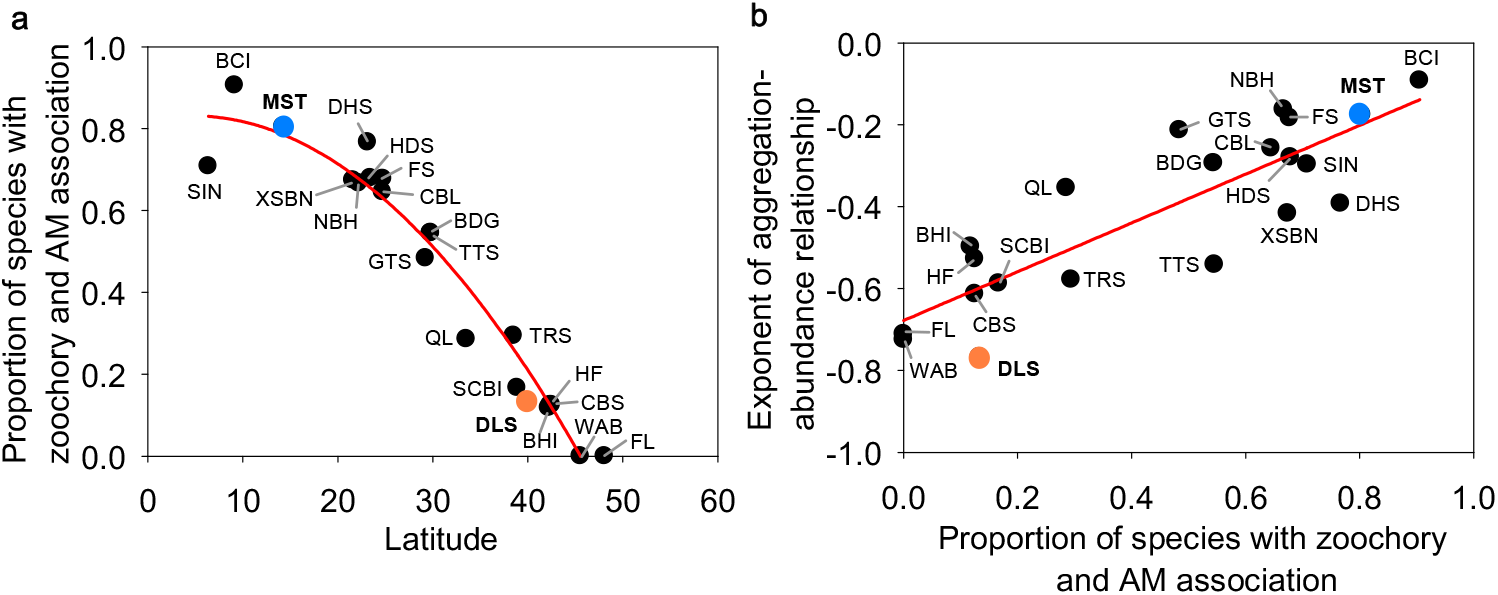
Latitudinal variation in the proportion of species showing mainly animal seed dispersal and arbuscular mycorrhizal (AM) association. **a**, latitudinal gradient in the proportion of species per plot that show both, mostly animal seed dispersal and AM association, **b**, relationship between the exponent of the aggregation-abundance relationship and the proportion of species per plot that show mostly animal seed dispersal and AM association for the 21 forest plots. See Extended Data Fig. 3 for the individual relationships with animal seed dispersal and AM association. To outline the overall tendency in the data we fitted in a) a polynomial regression of order 2 and in b) a linear regression. Note that most species showed either arbuscular mycorrhizal (AM) or ectomycorrhizal (EM) associations. Colours indicate the example species of Figure 2.

A theory that describes how the observed response of conspecific aggregation to abundance influences species coexistence requires a dynamic population model that relates competition at the population level to the emerging spatial patterns at the scale of individual trees. Here, we derive such an approach^8^, which incorporates the critical information on spatial patterns at the scale of individual trees provided by the ForestGEO datasets. We exemplify our theory by a simple model, which assumes that survival of individual trees is reduced in areas of high local tree density, as described by neighbourhood crowding indices^36,37^ (Box 1), while reproduction is assumed to be density-independent^7^:

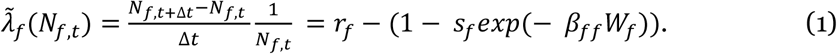

Here *N*_*f,t*_ is the abundance of the focal species *f* at time *t*; the time interval Δ*t* is the 5yr census interval; 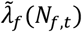 is the per-capita population growth rate (or average individual fitness^40^) of species *f* as function of species abundance *N*_*f,t*_; *s*_*f*_ is a density-independent per-capita background survival rate: *r*_*f*_ the per-capita recruitment rate, and *β*_*ff*_ the conspecific individual-level interaction coefficient. *W*_*f*_, the fitness factor^40^, is the average of the sum *C*_*kf*_ + *I*_*kf*_ of con- and heterospecific neighbourhood crowding taken over all individuals *k* of species *f* (Box 1, eq. d) and depends on the abundances of all species, their spatial patterns and the aggregation-abundance relationship (equations 7b, f, 9b).

Our spatially enriched macroscale model can be parameterized with the detailed information on spatial patterns extracted from forest megaplots (see methods). By taking a mean-field approach^41^ (i.e., diffuse competition at the community scale^8^) and assuming zero-sum dynamics^2^ (due to strong local density dependence), we can decouple the multispecies community model into multiple singe-species models (equation 7), where the multispecies fitness factor *W*_*f*_ of equation 7b is replaced by the single-species fitness factor (equation 7e).

The deterministic invasion criterion^1,5^ could be used to assess the consequences of the observed responses of conspecific aggregation to abundance on species coexistence. It asks whether a species will increase in abundance when it invades the resident community at low abundance, but ignores demographic stochasticity^14^ and species aggregation^9^. To overcome this limitation, we propose the plot-level extinction risk of species at low abundances as novel stochastic invasion criterion. To this end we translate the resulting single-species macroscale population model (equation 9) into a stochastic birth-death model^42,43^ that uses deterministic equations (equations 12, 13) to describe the dynamics of the probability *P*_*n*_(*t, N*_0_) that a species with initially *N*_0_ individuals has abundance *n* at timestep *t*. In particular, we are interested in the probability of non-successful invasion, given by the probability *P*_0_(*t, N*_0_) that a species starting with a small abundance *N*_o_ is extinct at timestep *t*. Although the time-dependent extinction probability *P*_0_(*t, N*_0_) of the birth-death model (equations 12, 13) can only be solved in special cases^43^, we derived a good approximation for our model (Fig. 4a; Extended Data Figs. 4, 5)

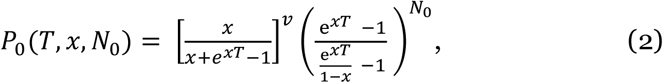

with 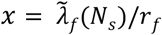 and *T* = *r*_*f*_*t*, where 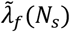 is the per-capita population growth rate of species *f* (equation 9) at a suitable small abundance *N*_s_, *r*_*f*_ the recruitment rate, *T* the scaled time step, and the parameter *v* determines the immigration rate (which is *ν r*_*f*_; equation 12a). Thus, the extinction dynamics of the birth-death model for small initial abundances *N*_o_ approximate that of a linear birth-death model^43^ where the birth and death rates (equations 12a, b) are taken at a typical abundance *N*_*s*_ (inset Fig. 4a; Extended Data Figs. 4). This equation provides the desired quantitative link between the extinction risk of a species when rare (i.e., our stochastic invasion criterion) with demographic rates, spatial patterns and immigration of species.

**Fig. 4.**
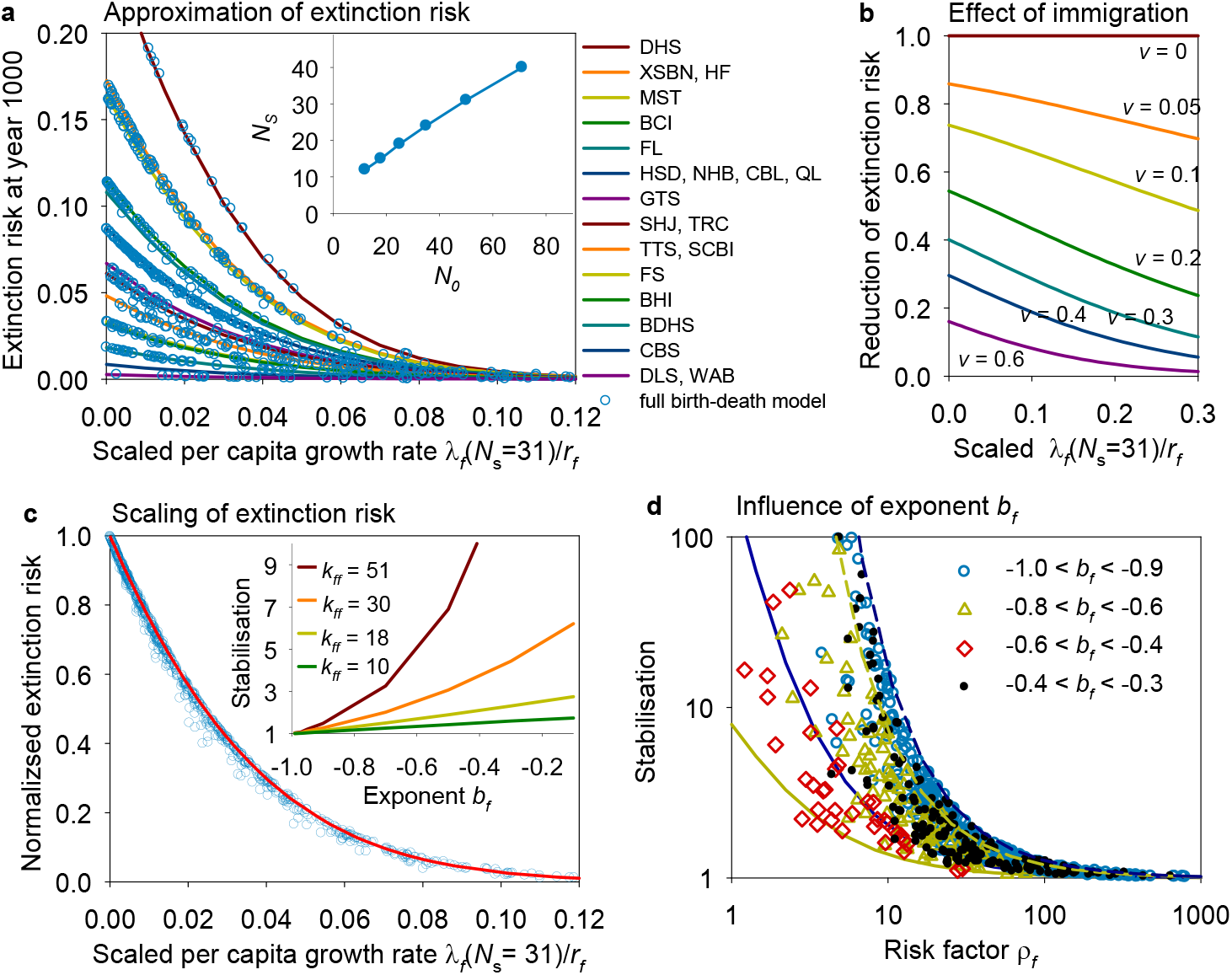
The stochastic invasion criterion. **a**, shows the plot-level extinction risk at year 1000 (i.e., *t* = 200) as a function of the scaled per-capita population growth 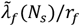 and for different forest plots. All species start with *N*_0_ = 50 individuals. The lines show the approximation with equation 2 and the open disks are the results of the numerical iteration of the birth-death model for the different species of the 21 forest plots. The inset shows how the fitted typical abundance *N*_*s*_ depends on the initial abundance *N*_0_, **b**, shows how immigration reduces the extinction risk, as given by the first term in equation 2, for a reproduction rate of *r*_*f*_ = 0.1. The constant immigration rate is given by *r*_*f*_*v*, and a value of *v* = 0.1 means one immigrant every 100 time steps on average. **c**, same as a), but we normalized the extinction probability with that of the corresponding neutral model. The small inset shows how stabilisation (i.e., the invers of the normalised extinction risk) depends for small observed abundances 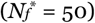 on aggregation *k*_*ff*_ and the exponent *b*_*f*_ of the aggregation-abundance relationship, estimated with equation 2 for parameters *k*_*fh*_ = 0.95, *J*^*^ = 21000, *B*_*f*_ =1, *N*_o_ = 80, *r*_*f*_ = 0.1, and *v* = 0. **d**, Stabilization, the factor by which non-neutral mechanisms reduce the extinction risk, for the 720 species of the 21 forest plots, plotted over the risk factor *ρ*_*f*_ = 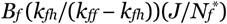 (equation 16c). Different values of the exponent *b*_*f*_ are shown by coloured symbols. Additionally, we show for different typical cases how stabilisation depends on the risk factor *ρ*_*f*_: the gold lines are for *b*_*f*_ = −0.9 and the blue lines for *b*_*f*_ = −0.3, dashed lines indicate 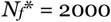 and solid lines 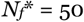. The data in a), c), and d) are from scenario 1 (i.e., no niche differences, *N*_o_ = 50, no immigration).

To assess the effects of spatial structure and other non-neutral mechanisms on the invasion success of species when rare, we normalized the extinction risk *P*_0_(*T, N*_0_) by the extinction risk of the corresponding neutral model without immigration (equation 15c). This allows us to quantify the degree of stabilisation, i.e., the factor by which the extinction probability declines due to spatial structure, niche differences and immigration. The scaled per-capita growth rate 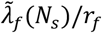, and thereby stabilisation (Fig. 4c), are determined by the observed species abundance 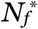, the exponent *b*_*f*_ of the aggregation-abundance relationship, and a risk factor given by 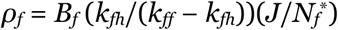 (equation 16). First, stabilisation strongly increases if *ρ*_*f*_ becomes smaller (Fig. 4d), which is the case if niche differences increase (i.e., *B*_*f*_ becomes smaller) and if the average neighbourhoods contain more conspecifics and less heterospecifics (i.e., *k*_*fh*_/(*k*_*ff*_ *− k*_*fh*_) becomes smaller).

Second, stabilisation decreases if the exponent *b*_*f*_ approaches values of −1 (i.e., stronger negative correlations between aggregation and abundance) (small inset Fig. 4c). For example, the solid gold line in Fig. 4d represents rare species at temperate forests 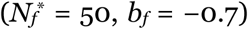 and the solid blue line rare species at tropical forests 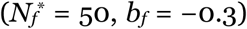. Finally, stabilisation increases with increasing abundance 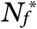 of the species, which is also included in the risk factor (as shown by comparison of solid versus dashed lines in Fig. 4d).

We find that coexistence in the analysed forest megaplots generally benefits from the emerging spatial patterns (Table 1a). For example, model results for scenarios without niche differences and without immigration indicate that the spatial patterns reduce the 1000-year local extinction risk of invading species for tropical, subtropical and temperate forests on average by factors 1.7, 1.9, and 2.7, respectively, with only moderate declines over longer time periods. Somewhat surprisingly, species at temperate forests tend to show on average higher stabilisation compared to tropical forests (Table 1a).

**Table 1.** Stabilization (the factor by which the local extinction risk is reduced) by different non-neutral mechanisms through time. We show the effect of combinations of different mechanisms on stabilisation, separately for all species in tropical, subtropical and temperate forests for our 4 scenarios. **a**, scenario 1 (no immigration: *v* = 0, no niche differences: *β*_*fi*_ = *β*_*ff*_), **b**, scenario 2: (no immigration: *v* = 0, niche differences: *β*_*fi*_ < *β*_*ff*_), **c**, scenario 3 (immigration with *v* =0.1, *β*_*fi*_ = *β*_*ff*_), and **d**, scenario 4 (immigration with *v* =0.1, *β*_*fi*_ < *β*_*ff*_). See Extended Data Fig. 6 for latitudinal gradients in stabilisation and temporal trends.

|  | 1000<br>years | 5000<br>years | 10,000<br>years | 1000<br>years | 5000<br>years | 10,000<br>years |
| --- | --- | --- | --- | --- | --- | --- |
|  | **a No immigration, no niches** |  |  | **c Immigration, no niches** |  |  |
| all | 1.91 | 1.79 | 1.68 | 2.62 | 3.07 | 3.27 |
| tropical | 1.74 | 1.62 | 1.51 | 2.42 | 2.81 | 2.96 |
| subtropical | 1.93 | 1.83 | 1.73 | 2.62 | 3.08 | 3.30 |
| temperate | 2.67 | 2.51 | 2.39 | 3.62 | 4.35 | 4.85 |
|  | **b No immigration, but niches** |  |  | **d Immigration and niches** |  |  |
| all | 2.73 | 2.46 | 2.23 | 3.78 | 4.34 | 4.55 |
| tropical | 2.58 | 2.29 | 2.04 | 3.63 | 4.10 | 4.23 |
| subtropical | 2.50 | 2.32 | 2.14 | 3.41 | 3.99 | 4.22 |
| temperate | 4.44 | 4.04 | 3.70 | 6.03 | 7.19 | 7.86 |

Using expanded model versions, we find that adding niche differences (Table 1b) or a small immigration (*v* = 0.1) (Table 1c) lead to further increases in stabilisation, and immigration and niche differences together reduce the plot-level extinction risk in tropical, subtropical and temperate forests by factors 6.0, 7.2, and 7.9, respectively (Table 1d). Finally, larger time periods lead to similar values of stabilisation (Table 1).

## Discussion

Developing an approach that integrates observed spatial patterns in forests with multiple ecological processes into mathematical theory is a considerable challenge. Here we present a unified framework integrating spatial point process theory with population models and their corresponding stochastic birth-death models to derive expectations for how interactions between multiple spatial patterns and processes impact stabilisation and species coexistence. The framework relies on spatial patterns that link competition at the scale of individual trees with the dynamics at the population scale. Our theory leads to a closed formula that shows how the extinction risk of species when rare depends on spatial patterns, demographic parameters, and immigration.

Several important ecological insights result from this new theory. First, we have shown that the spatial patterns found in forest megaplots have in general a stabilizing effect under neighbourhood competition^7,8^, as they substantially reduce the plot-level extinction risk of a species at low abundances. This result challenges a prevalent perspective^44,45^ that spatial patterns alone cannot foster coexistence. However, this assertion arises as a consequence of the assumption of previous spatially-explicit models^7,26,35,44,46^ of placing recruits close to their parents, which leads to destabilizing negative aggregation-abundance relationships (with exponents close to −1; inset Fig. 4c) where individuals of rare and abundant species experience similar degrees of conspecific competition (Fig. 1). Our results highlight the critical role that the aggregation-abundance relationship can play in shaping coexistence outcomes. Moving forward we suggest that models studying the forces structuring forest diversity and composition should include a more nuanced representation of seed dispersal^10,47^ and its interactions with mycorrhizal associations^11,12,13^.

Second, the observed latitudinal gradient suggests stronger stabilizing conspecific negative density dependence in tropical compared to temperate forests^13,22,48^ (Extended Data Fig. 7a, b). However, we found a general tendency for forests at lower latitudes to benefit less from the spatial coexistence mechanism (Table 1). This apparent contradiction can be explained by additional factors with opposed latitudinal tendencies that influence the extinction risk in our model (equation 17), such as lower risk factors *ρ*_*f*_ and higher species frequencies at higher latitudes (Extended Data Fig. 8). Nevertheless, species in temperate forests with a low equilibrium abundance (e.g., 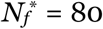) show a substantially lower stabilisation than species at tropical forests (small inset Fig. 4c) and therefore a higher extinction risk. This finding can help to explain why there are fewer rare species in temperate compared to tropical forests.

Third, we found that the frequently used deterministic invasion criterion^1,5^ did not yield useful predictions because it was satisfied for nearly all forests analysed here (equation 11) and failed to detect differences in the local extinction risk of species related to demographic stochasticity^14^. Additionally, it does not apply if rare species are more strongly aggregated than abundant species^9^, as observed here in temperate forests. This leads to conspecific population-level interaction coefficients that are not constant as commonly assumed^49^, but depend negatively on abundance (equation 7c). We have developed new theory that overcomes such issues and allows to determine whether and under what circumstances the abundances of rare species are likely to increase (equations 2, 16). Although we derived our stochastic invasion criterion for a specific population model, we believe that it could also be used for other population models that can be formulated by means of the per-capita growth rate 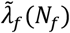 (see Supplementary Text).

Our spatial analysis of 21 large forest plots revealed an intriguing latitudinal gradient in the strength of the relationship between conspecific aggregation and abundance that begs for an explanation. Our data (Fig. 3) suggest that the aggregation-abundance relationship arises as an interaction of animal seed dispersal^10^ with mycorrhizal associations^11,12,13^. We hypothesize that an ectomycorrhizal (EM) association, as mostly observed in temperate forests, would be beneficial for species with dispersal close to adults (i.e., non-animal seed dispersal) because this allows facilitation of recruitment by ectomycorrhizal (EM) fungi near adults^11,12,13,39^. In contrast, animal seed dispersal farther away from conspecific adults is beneficial because the prevailing AM mycorrhizal associations in tropical forests offer less protection from enemies compared to an EM association^12,39^. Our combined dispersal-mycorrhiza hypothesis therefore provides an extra dimension to the study of negative conspecific density dependence, and an integrated understanding of the interacting effects of both, animal seed dispersal and mycorrhizal associations will be fundamental to our understanding of forces structuring forest diversity and composition.

Overcoming the limitations of macroscale population models, as often used in contemporary coexistence theory^1,4,15,40,50^, requires approaches that explicitly consider the lower-level processes that generate the phenomenon of interest^51^. Our results demonstrate the utility of transfer functions that describe the parameters of the macroscale population models, here the population-level interaction coefficients *α*_*fi*_ (between species *i* and *f*), as a function of individual-level interaction coefficients *β*_*fi*_ and measures of the emerging spatial patterns (i.e., *k*_*ff*_, *k*_*fh*_, *B*_*f*_; equation 7)^8^. Our scaling approach has the decisive advantage that the lower-level parameters of the transfer functions can be parameterized from empirical or experimental knowledge, such as the ForestGEO megaplots^6^ in our case. Taking this approach, we have demonstrated that spatial patterns emerging from individual-level processes play a key role in species coexistence, which underscores the need to understand the mechanisms underlying the spatial heterogeneity of forests in greater detail.

## Methods

### Study areas

Twenty-one large forest dynamics plots of areas between 20 and 50 ha with similar number of tropical, subtropical, and temperate forests were utilized in the present study (Supplementary Table S1). The forest plots are part of the Forest Global Earth Observatory (ForestGEO) network^6^ and are located in Asia and the Americas ranging in latitudes from 6.40° N to 48.08° N. Tree species richness among these plots ranges from 36 to 468. All free-standing individuals with diameter at breast height (dbh) ≥1 cm were mapped, size measured, and identified. We focussed our analysis here on individuals with dbh ≥ 10 cm (resulting in 313,434 individuals) and tree species with more than 50 individuals (resulting in initially 737 species). The 10 cm size threshold excludes most of the saplings and enables comparisons with previous spatial analyses. Shrub species were excluded. We also excluded 15 species with low aggregation (i.e., *k*_*ff*_ < *k*_*fh*_; Box 1), which would lead to negative growth rates at small abundances; ten of them from BCI, two from MST, two from NBH, and one from FS. These (generally less abundant) species are probable relicts of an earlier successional episode when they were more abundant^18,52^. We also excluded the two species *Picea mariana* and *Thuja occidentalis* of the Wabikon forest that are restricted to a patch of successional forest that was logged approximately 40 years ago^53^.

Most forest plots (18 of our 21 plots, those with more than one census) allow for estimation of the average mortality risk of individuals with dhh ≥ 10cm within one census period. We estimated mortality across all species and obtained for each forest plot one average mortality rate for trees with dbh ≥10 cm (Supplementary Table 1). In contrast, direct estimation of the per-capita recruitment rate is difficult. We therefore used a shortcut and assumed approximate equilibrium where the unknown per-capita recruitment rate *r*_*f*_ is the same as the known per-capita mortality rate. However, the equation of the local extinction probability (equation 2) we derived for the singe-species model (equation 9) allows for an easy assessment of the effects of species-specific recruitment rates. We also determined for all species used in our analyses the mycorrhizal association types based on available global datasets^54,55,56^ and website sources (http://mycorrrhizas.info/infex.html). To determine if a species is mainly dispersed by animals (zoochory) we used the Seed Information Database (https://data.kew.org/sid) of the Royal Botanic Gardens Kew and available literature^57^. Species without descriptions of mycorrhizal associations and dispersal modes were assigned according to their congeneric species. The proportion of focal species with zoochory, with arbuscular mycorrhizal (AM) association, and with both are shown in Supplementary Table 1.

### Proxy for pairwise competition strength between species

Some of our analyses require the ratio *β*_*fj*_/*β*_*ff*_ (Box 1) that describes the relative competitive effect^36,58^ of individuals of species *i* on an individual of the focal species *f*. In general, it is challenging to derive estimates for the pairwise interaction coefficients^37^ because this would require unfeasible large data sets to obtain a sufficient number of neighboured *f*–*j* species pairs for less abundant species. We therefore compared two scenarios. In scenario 1 we assumed that con- and heterospecific individuals compete equally, thus *β*_*fj*_ = *β*_*ff*_. In scenario 2 we assumed that individuals which are close relatives compete more strongly or share more natural enemies than distant relatives^59^. As proxy for this effect, we used phylogenetic distances^60^, given in millions of years (MA), as a surrogate for the relative competition strength because they are available for the species in our plots based on molecular data or the Phylomatic informatics tool^61^.

For plots without molecular data, we used the V. PhyloMaker2 package^62^ to generate a phylogenetic tree for each plot using GBOTB.extented.WP.tre updated from the dated megaphylogeny GBOTB^63^ as a backbone. For the other eight plots with molecular data, we followed the method reported by Kress et al.^64^ to build the phylogenetic tree based on DNA barcode data. We then used the cophenetic function in the picante package^65^ to calculate phylogenetic distance for each plot. In this we assume that functional traits are phylogenetically conserved^36,59,66^. To obtain consistent measures among forest plots, phylogenetic similarities were scaled between 0 and 1, with conspecifics set to 1 and a similarity of 0 was assumed for a phylogenetic distance of 1200 MA, which was somewhat larger than the maximal observed distance (1059 MA). This was necessary to avoid discounting crowding effects from the most distantly related neighbours^59^.

### Crowding indices for tree competition and measures of spatial patterns

We assume in our example model that survival of a focal tree *k* is reduced in areas of high local density of con- and heterospecifics (i.e., neighbourhood crowding), e.g., through competition for space, light or nutrients, or natural enemies^16,17,38,67^, while reproduction is density-independent with per-capita rate *r*_*f*_. Analogous models can be derived for crowding effects on the reproductive rate and/or the establishment of offspring (see Supplementary Text). We describe the neighbourhood crowding around tree *k* of a focal species *f* by commonly used neighbourhood crowding indices^36,37,58,68,69,70^ (*NCI*’s; Box 1), but use separate indices for con- and heterospecific trees.

The conspecific crowding index *C*_*kf*_ of a given individual *k* counts the number of conspecific neighbours *j* with distances *d*_*kj*_ smaller than a given neighbourhood radius *r*, but weights them by 1/*d*_*kj*_, assuming that farther away neighbours compete less (equation 3a). The heterospecific crowding index *H*_*kf*_ does the same with all heterospecifics (equation 3b), and the interaction crowding index *I*_*kf*_ weights heterospecifics additionally by their relative competitive strength *β*_*fj*_/*β*_*ff*_ (equation 3c). Thus, we estimate for each individual *k* three crowding indices

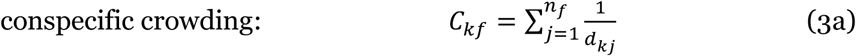

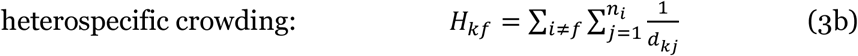

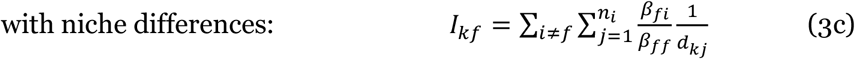

where *n*_*i*_ is the number of neighbours of species *i* within distance *r* of the focal individual, *d*_*kj*_ is the distance between the focal individual *k* and its *j*th neighbour of species *i*, and *β*_*fi*_/*β*_*ff*_ is the competitive effect of one individual of species *i* relative to that of the focal species *f* ^36,37,59,68,69,70^.

To link the survival of an individual *k* to its crowding indices we follow earlier work on individual neighbourhood models^36,37,59,69,70^ and assume that the survival rate *s*_*kf*_ of a focal tree *k* of species *f* is given by

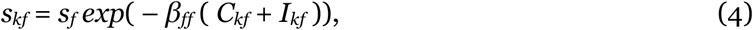

where *s*_*f*_ is a density-independent background survival rate of species *f* and *β*_*ff*_ the individual-level conspecific interaction coefficients of species *f*^36^. Statistical analyses with neighbourhood crowding indices have shown that the performance of trees depends on their neighbours mostly within distances *r* of up to 10 or 15 m^70^, we therefore estimate all measures of spatial neighbourhood patterns with a neighbourhood radius of *r* = 15 m. We investigate two scenarios, in scenario 1 con- and heterospecifics competing equally at the individual scale (i.e., *β*_*fi*_ = *β*_*ff*_), and in scenario 2 the quantity *β*_*fi*_/*β*_*ff*_ is proportional to phylogenetic similarity see above “*Proxy for pairwise competition strength between species”*).

We use scale transition theory^40^ and spatial point process theory^25^ to transfer the individual-based microscale information on the number and distance of con- and heterospecific neighbours of focal individuals, which are provided by the Forest-GEO census maps, into macroscale models of community dynamics^8^. To this end, we average the survival rates *s*_*kf*_ of all individuals *k* of the focal species *f* to obtain the average population-level survival rate 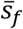, for which we derived a closed expression for Gamma distributed crowing indices^8^:

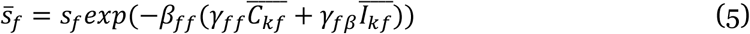

where 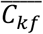 and 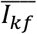 are the average crowding indices, and the quantities *γ*_*ff*_ and *γ*_*fβ*_ arise through the averaging because of the non-linearity in equation 4^8^, but in our case of high survival rates they are near one and can be neglected (i.e., *γ*_*ff*_ ≈ 1 and *γ*_*fβ*_ ≈ 1).

To incorporate the average survival (equation 5) into our population model we need to decompose the average crowding indices into species abundances and measures of spatial patterns (Box 1). Briefly, we do this by expressing the crowding indices in terms of the pair correlation function, a basic summary function of spatial statistics^25^, and the mean density *λ*_*f*_ = *N*_*f*_/*A* of the species *f* across the whole plot of area *A* (see equations S1-S8 in Supplementary Text). The resulting measures *k*_*ff*_ and *k*_*fh*_ of spatial patterns quantify the change in average conspecific and heterospecific neighbourhood crowding, respectively, relative to the case without spatial patterns (i.e., random distribution of the focal species and independent placement of the focal species with respect to the heterospecifics^25^):

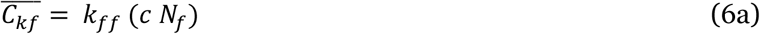

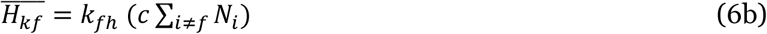

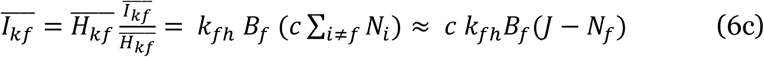

where *c* = 2 π *r*/*A* is a scaling factor (see equation S7 in Supplementary Text), *A* the area of the plot, *r* the radius of the neighbourhood, *J* the total number of individuals in the plot, and 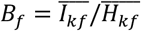 the average competition strength of one heterospecific neighbour relative to that of one conspecific. The quantity *k*_*ff*_ measures spatial patterns in conspecific crowding of species *f* (*k*_*ff*_ > 1 indicates aggregation, and *k*_*ff*_ < 1 regularity), and *k*_*fh*_ measures patterns in heterospecific crowding around the focal species *f* (*k*_*fh*_ < 1 indicates segregation, and *k*_*fh*_ > 1 attraction). Note that our measure of conspecific aggregation, which weights neighbours by distance, is correlated to Condit’s Omega measure of aggregation^19^ that counts the number of neighbours without weighting by distance (Extended Data Fig. 9). Additionally, we found that the strength of the latitudinal gradient was for a radius of say *r* > 10m basically independent of the neighbourhood area over which conspecific aggregation was measured (Extended Data Fig. 2). This was expected because of the distance-weighting (Box 1) where distant neighbours contribute little to total neighbourhood crowding.

Equation (6c) uses two findings of earlier work. First, a crucial insight used in our approach^8^ is that crowding competition of individual trees, as described by equation 4, leads in species-rich communities to diffuse competition at the population scale. That is, when taking a mean-field approximation^41,71^, the species-specific competition strengths of heterospecifics can be replaced in the macroscale model by the average heterospecific competition strength *B*_f_ ^8^, which summarizes the emerging effects of the individual-level interaction coefficients *β*_*fi*_/*β*_*ff*_ at the population-level. For species rich forests at or near a stationary state, *B*_*f*_ is in good approximation a species-specific constant (see Supplementary Text in Wiegand et al.^8^). Second, the strong local density dependence causes approximate zero-sum dynamics, where the total number *J* of individuals remains constant^2^, and the number of heterospecifics is given by *J* − *N*_*f*_ (equation 6c). Using these approximations in equation 6c decouples the multispecies dynamics and allows us to investigate the dynamics of individual species in good approximation.

In Fig. 2 we fitted for the species of a given forest plot a phenomenological power law where the x-value was the logarithm of abundance *N*_*f*_ and the y-value the corresponding logarithm of *k*_*ff*_ ^20,27,28^. However, this leads for large abundances to values of *k*_*ff*_ close to zero which would indicate strongly regular patterns^25^ not found in the data. Instead, in the extreme case without an aggregation mechanism (i.e., random placement of offspring) crowding competition leads to repulsion of conspecifics comparable to segregation *k*_*fh*_ of heterospecifics. Thus, to avoid a bias we used the quantity *k*_*ff*_ − *k*_*fh*_ as the y-value in our fit.

### Basic community-level model

Combining equations (1), (5) and (6) leads to the spatially-enriched macroscale model for the per-capita growth rate of species *f* (also called average individual fitness^40^):

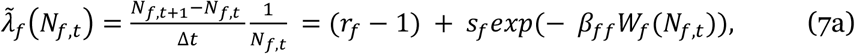

where *N*_*f,t*_ is the abundance of species *f* at timestep *t, s*_*f*_ is a density-independent per-capita background survival rate, *r*_*f*_ the per-capita recruitment rate, *β*_*ff*_ the individual-level conspecific interaction coefficients of species *f*, and *W*_*f*_(*N*_*f,t*_) the fitness factor^40^ given by

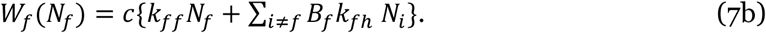

The community-level interaction coefficients are therefore given by:

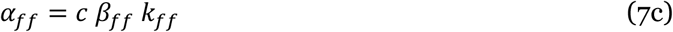

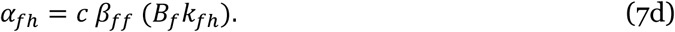

Thus, even if the individual-level interaction coefficients *β*_*fi*_ (equations 3a, c) are constant, the community-level conspecific interaction coefficients *α*_*ff*_ (equation 7c) are not necessarily constant, as is commonly assumed^49^. Instead, they will depend on abundance if aggregation *k*_*ff*_ correlates with abundance^8^, as observed in many of our forest dynamics plots (Fig. 2). Assuming approximate zero-sum dynamics we can rewrite the fitness factor to

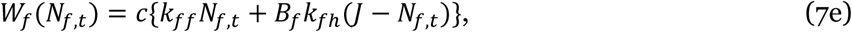

and to accommodate the power law in (*k*_*ff*_ - *k*_*fh*_) we further rewrite the fitness factor to:

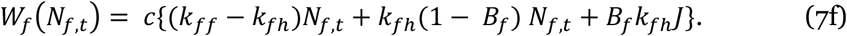

Our spatially-enriched macroscale model approximates an underlying individual-based model in the tradition of earlier spatially-explicit work^7,35,45,46,72,73^, but uses the empirically observed spatial patterns instead of modelling their dynamics explicitely^8^. To parameterize the model, we used the ForestGEO data sets to determine the values of aggregation *k*_*ff*_, segregation *k*_*fh*_, total community size *J*, the observed abundance 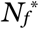 of species *f*, the recruitment rate *r*_*f*_, and the exponent *b*_*f*_ that describes the scaling of aggregation with abundance (equation 8). In all analyses we set the background survival rate to *s*_*f*_ = 1 (i.e., a tree without neighbours within distance *r* will survive this timestep). To determine the unknown value of *β*_*ff*_, the individual-level conspecific interaction coefficients of species *f*, we assume that the observed species abundance *N*_*f*_ is close to equilibrium 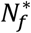 and obtain from equation 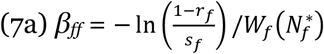. The influence of uncertainty in the equilibrium abundance 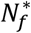 on our results can be assessed from equations 16 and 17. The effects of large uncertainty (e.g., 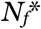 of 50 vs. 2000) is illustrated in Fig. 4d. Note however that the stochastic birth-death model derived from our macroscale model (that considers demographic stochasticity) describes for species with low values of the percapita growth rate 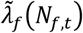 non-equilibrium behaviours where the abundances fluctuate (together with aggregation) stochastically.

To consider the observed aggregation-abundance relationships (see Extended Data Fig. 10 for plots) we replace in equation (7f) the quantity *k*_*ff*_ − *k*_*fh*_ by

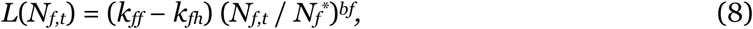

where 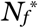 is the observed species abundance. Thus, 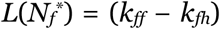. Note that effects of habitat association or details of dispersal will influence the values of *k*_*ff*_ and cause the observed departures from the aggregation-abundance relationship. With equation (8) we obtain our final spatially enriched model that links the macroscale to the microscale:

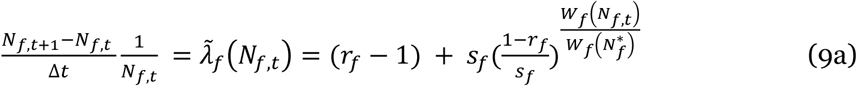

with

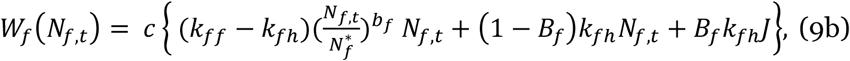

Note that we obtain for the case of *s*_*f*_ = 1 (i.e., all mortality depends on neighbourhood crowding) and for small per-capita recruitment rates *r*_*f*_ the approximation

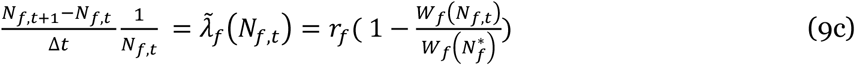

which suggests using a scaled mean recruitment rate 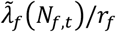 to remove the effect of the recruitment rate.

### The deterministic invasion criterion

Coexistence requires that populations at low abundances *N*_*f*_ increase^1,5^, thus 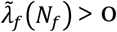. This is the case if 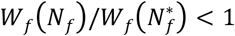 (equations 9a, c), which leads to:

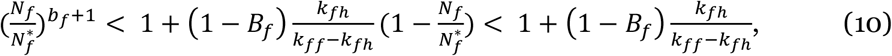

and with 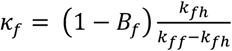 and *N*_*f*_ = 1 we find the deterministic invasion criterion

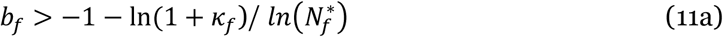

In the extreme case that con- and heterospecifics competing equally (i.e., *B*_*f*_ = 1) we find:

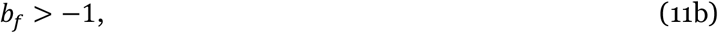

which is always fulfilled if the exponent *b*_*f*_ > -1. In the more realistic case where heterospecifics compete weaker than conspecifics (i.e., *B*_*f*_ < 1), equation (11a) indicates that the invasion criterion can be fulfilled even if *b*_*f*_ < −1. Indeed, some species of the CBS plot, which shows an exponent of *b*_*f*_ = −1.077, fulfil the deterministic invasion criterion due to large values of *k*_*fh*_/(*k*_*ff*_ - *k*_*fh*_) (and *B*_*f*_ < 1; Extended Data Fig. 7e,f). An aggregation-abundance relationship with exponent *b*_*f*_ < −1 leads to a type of Allee effect, which reduces the growth rate at low abundances or makes it even negative (Extended Data Fig. 7).

### The stochastic invasion criterion

The deterministic invasion criterion ignores the detrimental influence of demographic stochasticity, which can produce a high risk of extinction at small population sizes, even if the per-capita growth rate 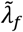 shows positive values^14^. To be able to consider the impact of demographic stochasticity and immigration, we translate 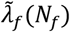 into the master-equation framework of a Markov process of birth-death type^42,43^, to estimate the probability distribution *P*_o_(*t, N*_o_) that a species with initially *N*_0_ individuals will be extinct at timestep *t*^74^. A convenient property of the birth-death model is that it uses deterministic equations to describe demographic stochasticity and does therefore not require stochastic simulations to determine extinction probabilities.

The birth-death model describes the dynamics of the probabilities *p*_*n*_(*t*) that the population at time *t* has abundance *n*. To that end, the model considers the rates *b*_*n*_ and *d*_*n*_ at which reproduction and death events occur. We find from equation (7a) that

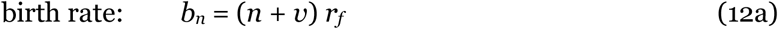

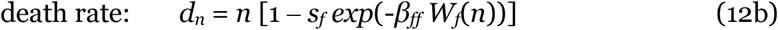

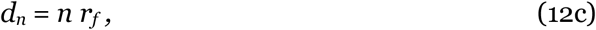

where the birth rate in equation (12a) includes immigration with a constant rate *v r*_*f*_, and equation (12c) gives the death rate of the corresponding neutral birth-death model. Note that we model immigration via a constant rate *ν r*_*f*_ ^42,43,75^. This approach differs from the way immigration is usually modelled in neutral theory^2,26,76^.

The birth-death model is given by the differential equation system

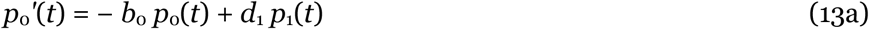

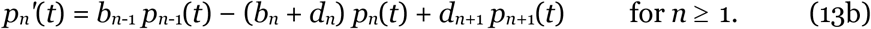

The first equation (13a) shows how the extinction probability *p*_o_(*t*) changes at time *t*. It increases if death events occur in populations with abundance *n* = 1 (happening at rate *d*_1_) and decreases if new individuals immigrate to extinct populations (happening at rate *b*_o_ = *v r*_*f*_). To obtain the effective change rate of *p*_o_(*t*), the rates *d*_1_ and *b*_o_ are multiplied with the corresponding probabilities *p*_1_(*t*) and *p*_o_(*t*) that the population has one or no individual, respectively. The second equation (equation 13b) keeps track of birth and death events in existent populations. Birth events occur in a population with *n*–1 individuals (which exists with probability *p*_*n*-1_(t)) at rate *b*_*n*-1_. Hence, these events increase the probability *p*_*n*_(*t*) of population size *n* at rate *b*_*n*-1_*p*_*n*-1_(*t*). Similarly, death events occur in populations of *n* + 1 individuals (probability *p*_*n+*1_(t)) at rate *d*_*n*+1_ and increase *p*_*n*_(*t*) at rate *d*_*n*+1_ *p*_*n*+1_(*t*). Conversely, for populations with *n* individuals, both birth and death events (occurring at rates *b*_*n*_ and *d*_*n*_, respectively) reduce the population probability *p*_*n*_(*t*), leading to a reduction of *p*_*n*_(*t*) at joint rate (*b*_*n*_ + *d*_*n*_) *p*_*n*_(*t*). We solved the system (13a)-(13b) with an implicit integration technique. Details can be found in the supplemental software repository.

For assessment of the invasion criterion we iterated equation (13) in a straight-forward way using a small abundance *n*_o_ as initial condition *p*_*n*_(*t* = 0). In most of our examples we used a small abundance of *n*_o_ = 50, but conducted also analyses for *n*_o_ = 12, 18, 25, 36, 50, and 71 (Fig. 4a). We also tested the birth-death model with a symmetric individual-based model (i.e., all species have the same parameters and follow the same rules) described in Wiegand et al.^8^ and the Supplementary Text. To this end we used the initial abundance distribution of the individual-based simulations also in the birth-death model. Our test showed that the mean-field approximation works; the birth-death model, parameterized by spatial patterns, was able to correctly predict the species abundance distribution of the individual-based simulations for the same initial conditions and simulation period (Fig. S1 in Supplementary Text).

Given that we are here only interested in the extinction probability *P*_o_(*t, N*_o_) for small initial abundances *N*_o_, we approximate the density-dependent death rate in equation (12b) by a constant *d*_*f*_, being the death rate 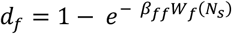 at a typical low abundance *N*_s_:

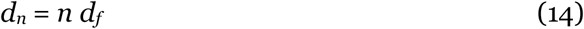

Now both, the birth rate *b*_*n*_ (equation 12a) and the death *d*_*n*_ are linear in *n* and the corresponding master equation (13) can be solved exactly^43^:

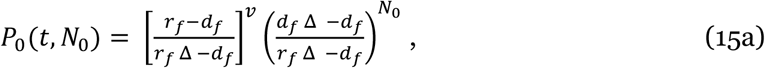

where 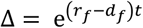. We note that *r*_*f*_ - *d*_*f*_ is the per-capita growth rate 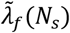 at the small abundance *N*_s_. We define 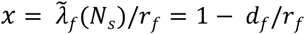 and *T* = *r*_*f*_*t* being the scaled time. With these definitions we obtain Δ = e^*xT*^ and therefore

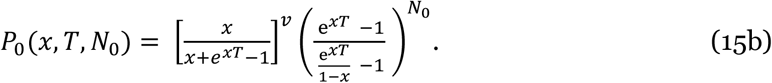

Thus, our approximation of the extinction risk is a function of the per-capita population growth rate at a small abundance divided by the recruitment rate *r*_*f*_ (i.e., the quantity *x*), but it depends additionally on the scaled time step *T* = *r*_*f*_ *t*, the initial population size *N*_o_, and the immigration parameter *v*. The extinction probability of the corresponding neutral model (where *x* ⟶ 0) is given by

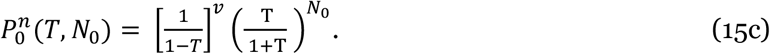

To apply the approximation of equation 15 to the results of the numerical iteration of the birth-death model (equations 12 and 13), we need to fit (for each value of *T* and *N*_o_) the value of the small abundance *N*_s_. This can be done in a straight-forward way by minimizing e.g., the sum of squares between the extinction risks predicted by the full birth-death model and its approximation for different values of *N*_s_ (Extended Data Fig. 4). Thus, the extinction dynamics of the birth-death model for small initial abundances *N*_o_ (and not too large timesteps *t*) approximates that of a linear birth-death model^43^, but with a per-capita growth rate of 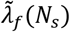 for a typical abundance *N*_*s*_ that is somewhat below the initial abundance *N*_o_ (Fig. 4a). By repeating this exercise for different initial abundances *N*_o_, ranging from 12 to 71, and different time periods of 1000, 5000, and 10000 years, we found still good fits and smooth functions for the best fitting abundance *N*_*s*_ (Fig. 4a; Extended Data Fig. 5)

### Link between extinction risk and species properties

The link between the plot-level extinction risk of small populations and the scaled average individual fitness 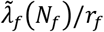 (equation 2) allows us to express the extinction risk as a function of spatial patterns (*k*_*ff*_, *k*_*fh*_, *B*_*f*_), demographic parameters (*r*_*f*_, *s*_*f*_) and the relative abundance of the species 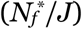. We note that the scaled average individual fitness at the typical abundance *N*_*s*_ is approximated by 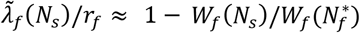 (equation 9c). We therefore investigate the ratio 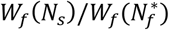 in more detail, it can be expressed as

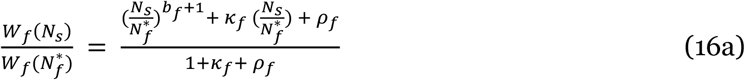

with

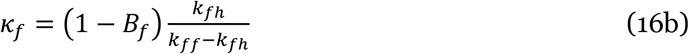

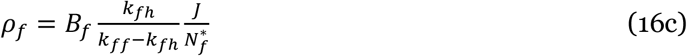

We can ignore in equation (16a) the quantity *k*_*f*_ if *B*_*f*_ (1 + *J/N*_*s*_) >> 1, which approximates *B*_*f*_ >> *N*_*s*_/*J* for small relative abundances *N*_*s*_/*J* of the focal species, and obtain

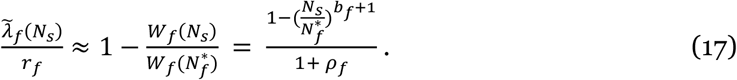

Thus, the scaled per-capita growth rate, and the reduction in the plot-scale extinction risk due to non-neutral mechanisms, is mostly driven by a risk factor *ρ*_*f*_ and modified by the term 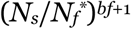 introduced by the abundance-aggregation relationship with exponent *b*_*f*_. It is therefore instructive to plot the factor by which the extinction risk of the corresponding neutral model is reduced (i.e., stabilisation) in dependence of the risk factor *ρ*_*f*_ to assess the relative impact of abundance 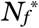 vs. the exponent *b*_*f*_ of the aggregation-abundance relationship (Fig. 4d). Interestingly, the risk factor *ρ*_*f*_ drops out in the deterministic invasion criterion of equation 10, which indicates that it misses important information.

### Scenarios investigated with the birth-death model

We conducted four experiments with the birth-death model to assess the effects of the spatial mechanism of neighbourhood crowding and immigration on the ability of species to persist for 1000 years, 5000 years and 10,000 years when having low abundances (i.e., *N*_o_ = 50). Scenarios 1 assumed that con- and heterospecifics compete equally (i.e., no niche differences; *β*_*fi*_ = *β*_*ff*_, *B*_*f*_ = 1), and scenarios 2 considers niche differences between species approximated by phylogenetic dissimilarity (see above “*Proxy for pairwise competition strength between species*”). Scenarios 3 and 4 are the same as scenarios 1 and 2, but additionally assume a small constant immigration with parameter *v* = 0.1. For the mean reproduction rate of *r*_*f*_ = 0.1 across all plots, this result in an immigration rate of *r*_*f*_ *v* = 0.01, or 1 immigrant every 100 timesteps. We also conducted analyses of scenario 1 with different initial conditions *N*_o_ = 12, 18, 25, 36, 50, and 71 to establish a relationship between *N*_o_ and the abundance *N*_*s*_ (Fig. 4a) that leads to the best fit of the extinction probability *P*_0_(*t, N*_0_) (derived by the numerical iteration of the full birth-death model) by our approximation of equation 2 (Supplementary Data Table 2a).

## Acknowledgments

X.W. was supported by the National Key Research and Development Program of China (2022YFF1300501), and the Key Research Program of Frontier Sciences, Chinese Academy of Sciences (Grant ZDBS-LY-DQC019). The Harvard ForestGEO Forest Dynamics plot was funded by the Center for Tropical Forest Science and Smithsonian Institute’s Forest Global Earth Observatory (CTFS-ForestGEO), the National Science Foundation’s LTER program (DEB 06-20443, DEB 12-37491, and DEB 18-32210) and Harvard University. Thanks to Jason Aylward and Christina McKeown for field supervision, data screening and database management. The FS plot project was supported by the Forestry and Nature Conservation Agency, the Taiwan Forestry Research Institute, and the National Science and Technology Council in Taiwan. Funding for the establishment of the SCBI ForestGEO Large Forest Dynamics Plot was provided by the Smithsonian Global Earth Observatory (SIGEO) initiative, the Smithsonian Institution National Zoological Park, and the HSBC Climate Partnership. The BCI censuses have been made possible through support of the U.S. National Science Foundation (awards 8206992, 8906869, 9405933, 9909947, 0948585 to S.P. Hubbell), the John D. and Catherine D. McArthur Foundation, and the Smithsonian Tropical Research Institute. Funding for the Tyson Research Center (TRC) Forest Dynamics Plot was provided by the National Science Foundation (DEB Awards 1557094 and 2240431 to J.A.M.), the International Center for Advanced Renewable Energy and Sustainability (I-CARES) at Washington University in St. Louis, Tyson Research Center, andForestGEO. We thank the Tyson Research Center staff for providing logistical support. The 25-ha Long-Term Ecological Research Project at Sinharaja World Heritage Site is a collaborative project of the Uva Wellassa University, University of Peradeniya, ForestGEO, with supplementary funding received from the John D. and Catherine T. Macarthur Foundation, the National Institute for Environmental Science, Japan, and the Helmholtz Centre for Environmental Research-UFZ, Germany, for past censuses. The PIs gratefully acknowledge the Forest Department, Uva Wellassa University, and the Post-Graduate Institute of Science at the University of Peradeniya, Sri Lanka for supporting this project. We thank D. Alonso and J. Capitan for discussion of birth-death models and the Kraft lab and R. Condit for valuable feedback on earlier drafts, and numerous local field and lab staff, technicians, interns, volunteers, and researchers for their invaluable contributions to the collection and management of the forest inventory data.

## Author contributions

T.W., X.W., S. F. and A. H. conceived and designed the project, T.W. implemented the models, conducted the simulations, analysed the results, and prepared figures and tables. T. W. and S. F prepared the software. T.W., A.H., S. F., X.W. and N.K. led the writing of the manuscript. X.W. assembled and analysed the plot data and conducted the spatial analyses. All other co-authors contributed data, reviewed, approved, and had the opportunity to comment on the manuscript.

## Extended Data Figures

**Extended Data Figure 1.**
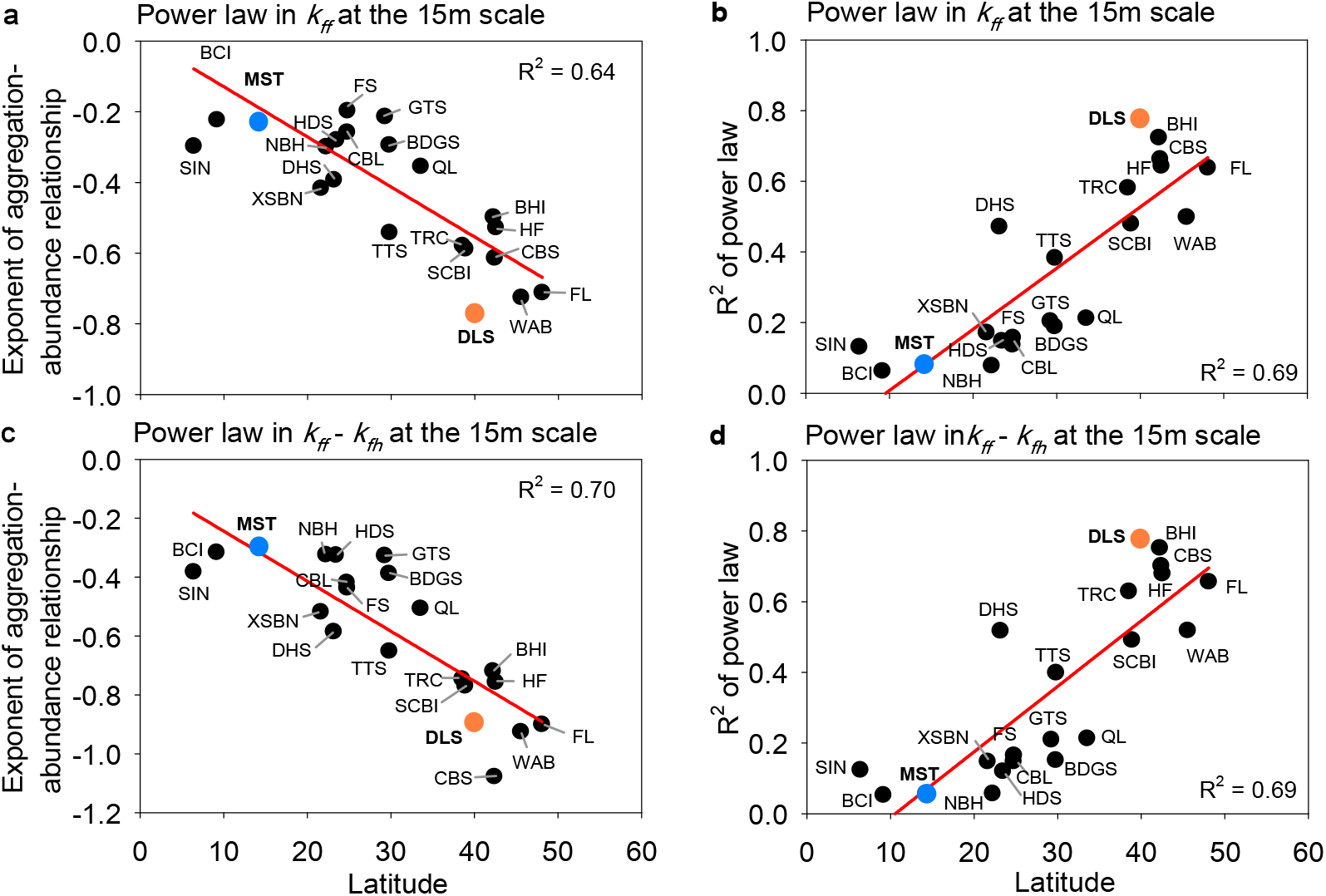
Latitudinal gradients in the slope of the abundance-aggregation relationship. **a**, the power law exponent of the scaling of aggregation *k*_*ff*_ with abundance *N*_*f*_, **b**, the *R*^2^ of the linear regression between ln(*k*_*fh*_) and ln(*N*_*f*_) for each species, **c**, same as a) but for the scaling of (*k*_*ff*_ - *k*_*fh*_) with abundance *N*_*f*_, **d**, same as b) but for the scaling of (*k*_*ff*_ − *k*_*fh*_) with abundance *N*_*f*_. Our measures of aggregation *k*_*ff*_ and segregation *k*_*fh*_ are based on neighbourhood crowding indices that count the number of neighbours within distance *r* of the focal individual, but each neighbour is weighted by the inverse of its distance to the focal individual (Box 1). The neighbourhood distance was in all panels *r* = 15m. For plot acronyms see Supplementary Table 1. For data see Supplementary Data Table 1. To show the overall tendency we fitted linear regressions.

**Extended Data Figure 2.**
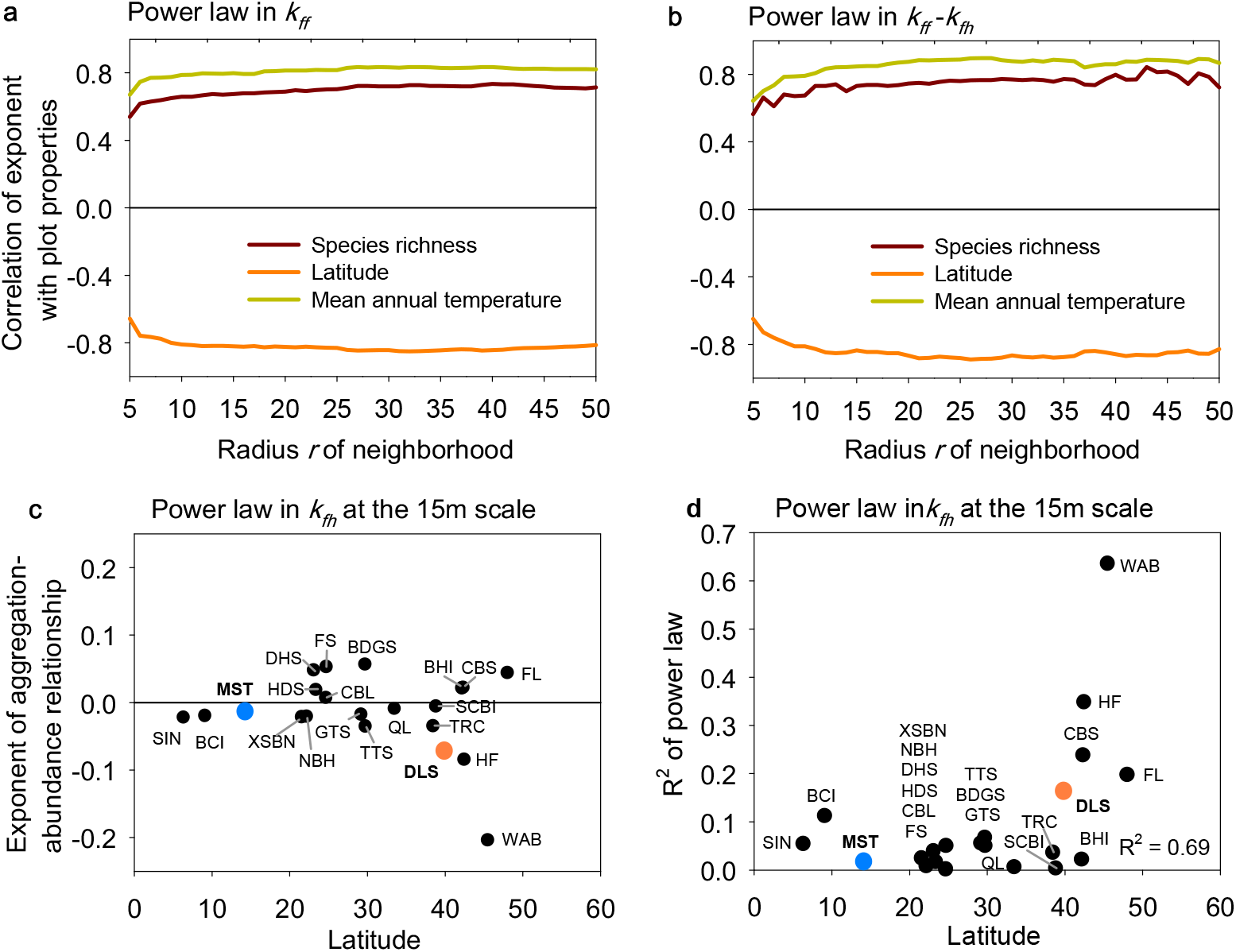
Abundance-dependency of aggregation *k*_*ff*_ and segregation *k*_*fh*_. **a**, the correlation coefficient between the exponent of the power law and the plot properties species richness, latitude and mean annual temperature for the power law in aggregation *k*_*ff*_ with respect to abundance *N*_*f*_, **b**, the power law in (*k*_*ff*_ – *f*_*fh*_) with respect to abundance *N*_*f*_, where *k*_*ff*_ is conspecific aggregation and *k*_*fh*_ heterospecific segregation, **c**, exponent of the power law of segregation *k*_*fh*_ with respect to species abundance *N*_*f*_ for the 21 the ForestGEO data sets, **d**, the *R*^2^ of the linear regression between ln(*k*_*fh*_) and ln(*N*_*f*_).

**Extended Data Figure 3.**
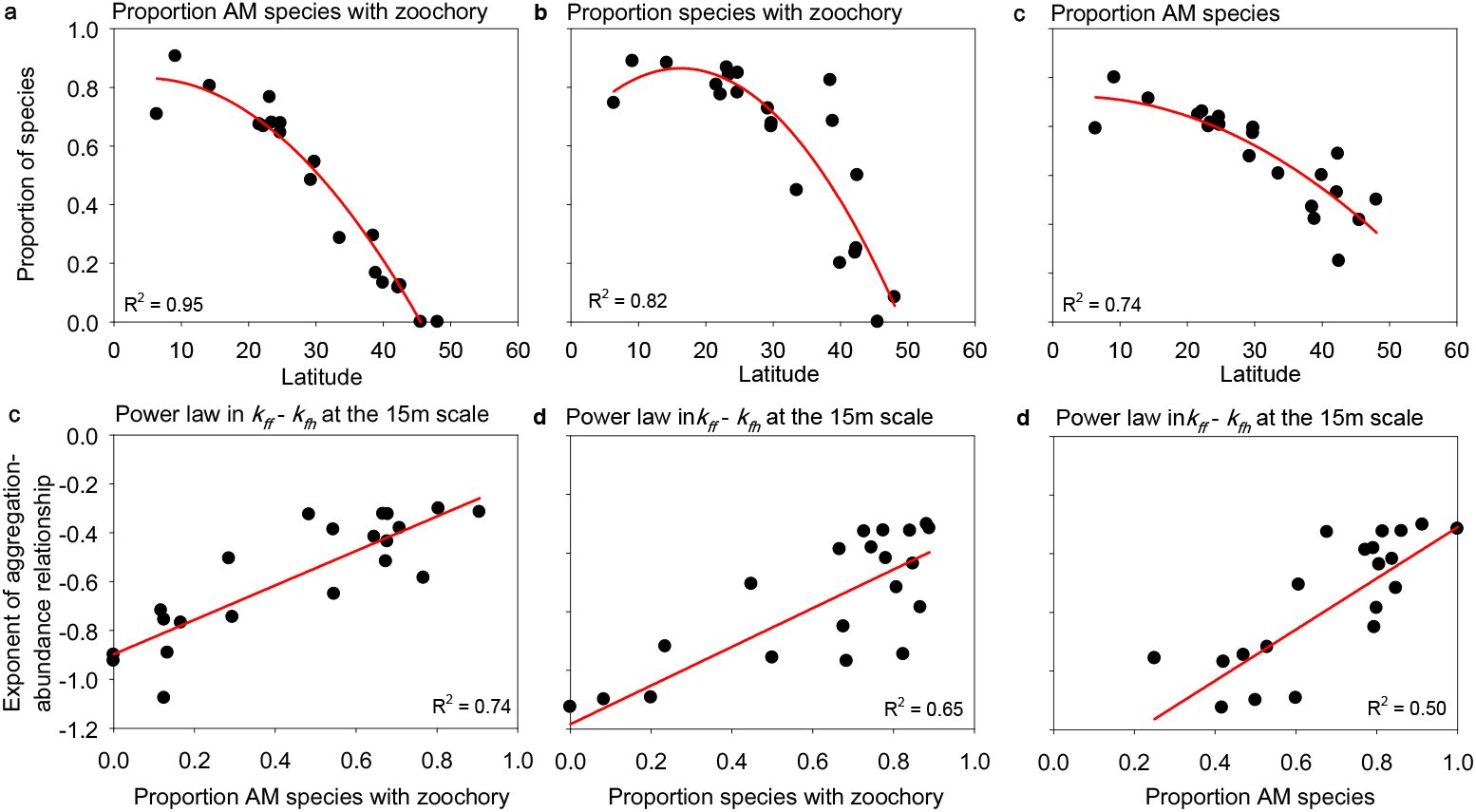
Latitudinal variation in the proportion of species showing zoochory and an arbuscular mycorrhizal (AM) association. **a**, latitudinal gradient in the proportion of species per plot that show both, an AM association and zoochory, **b**, same as a), but only zoochory, **c**, same as a), but only AM species, **d**, relationship between the slope of the aggregation-abundance relationship and the proportion of species per plot that show an AM association and zoochory, **e**, same as d), but only zoochory, **f**, same as d), but only AM species. To outline the overall tendency in the data we fitted in panels a) to c) a polynomial regression of order 2 and in panels d) to f) a linear regression.

**Extended Data Figure 4.**
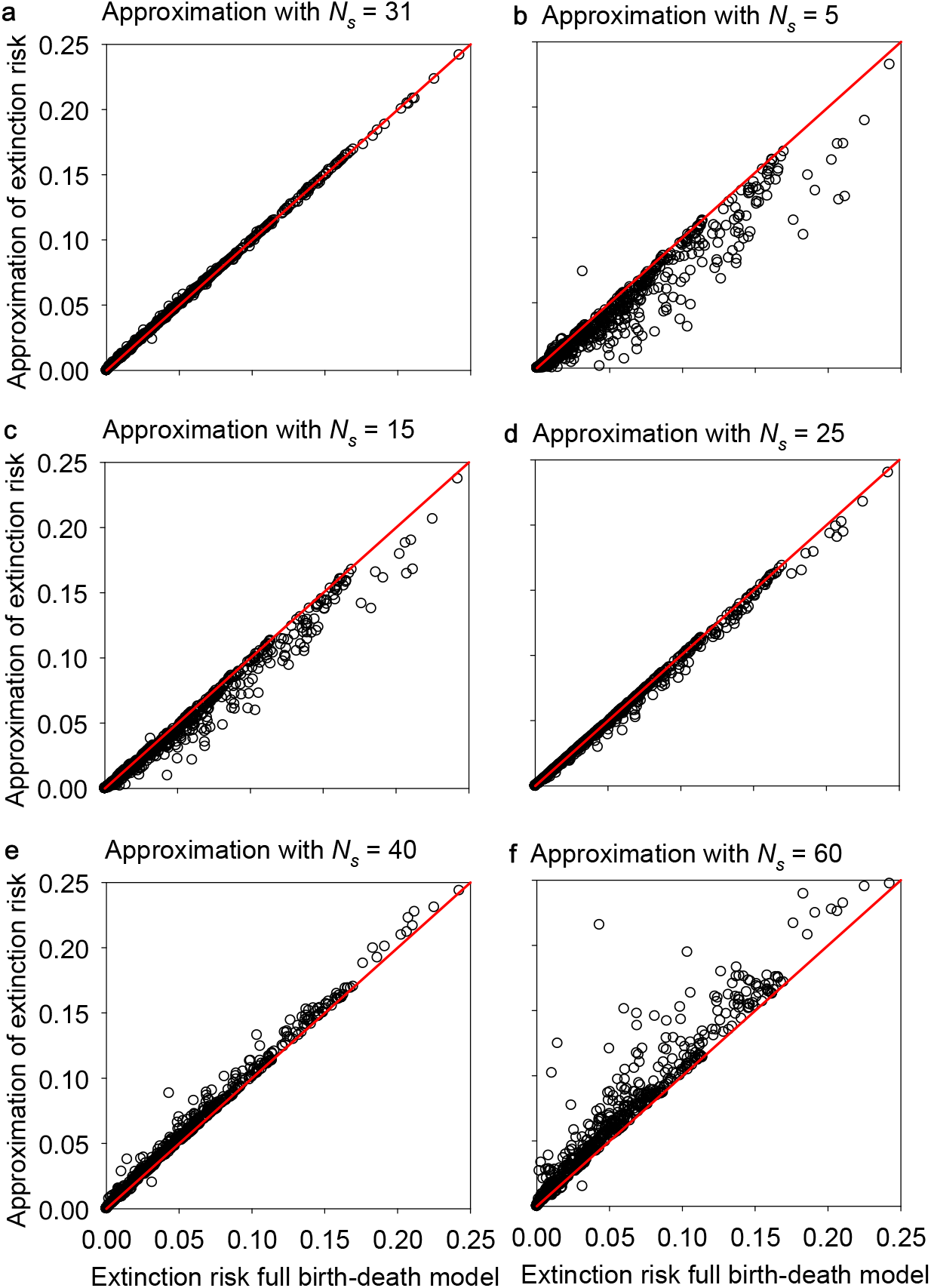
Approximation of the extinction probability of the birth-death model. The approximation with equation 2 requires determination of a typical abundance *N*_*s*_ to determine the constant mean population growth rate 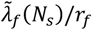. The different panels show the approximated extinction risk for different values of *N*_*s*_ over the extinction risk determined with the full birth-death model. Panel a) with *N*_*s*_ = 31 shows the best fit. The data are from scenarios 1 (no niche differences, *N*_o_ = 50, no immigration: *v* = 0), see Supplementary Data Table 1. The red line is the one-to-one line.

**Extended Data Figure 5.**
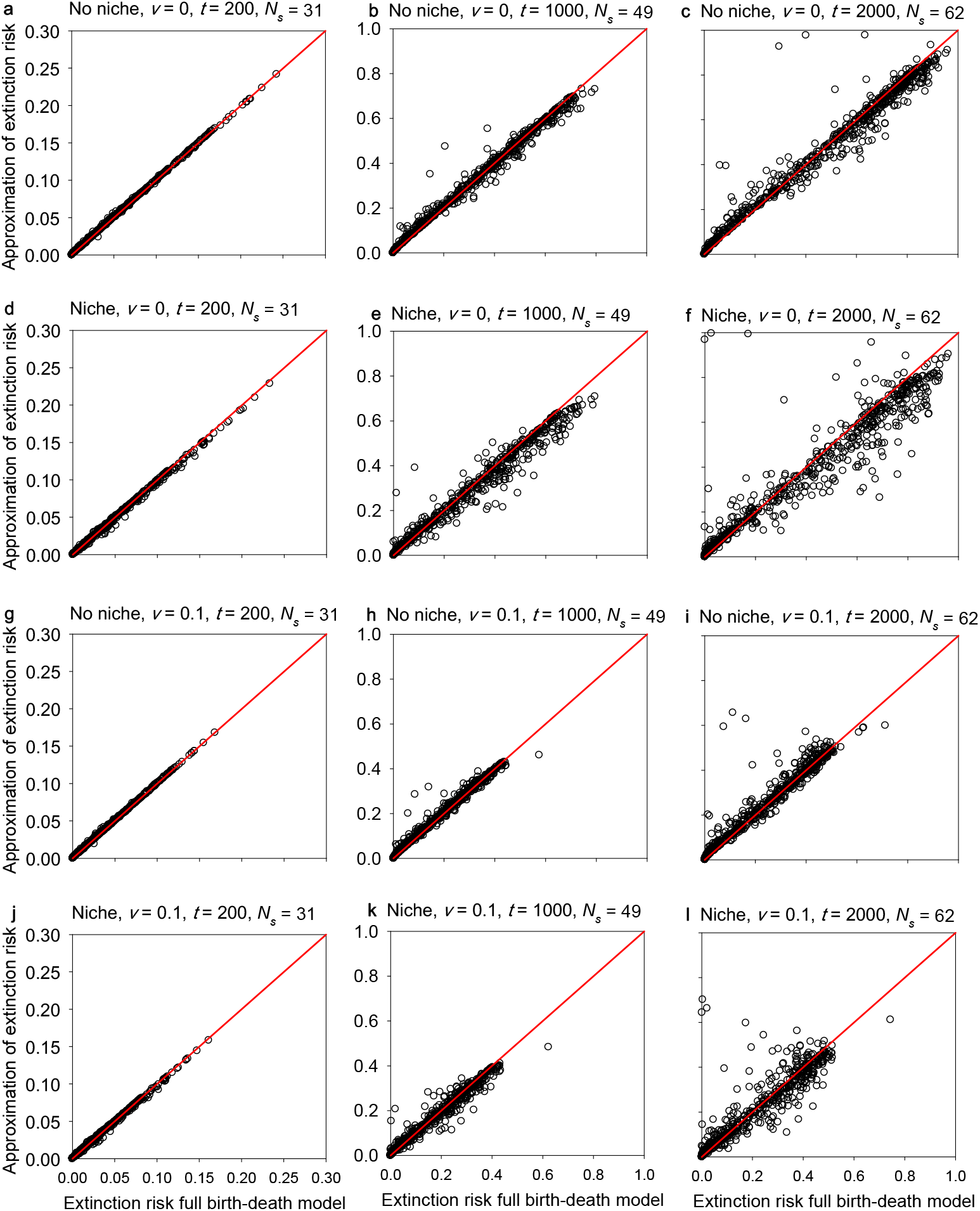
Approximation of the extinction probability of the birth-death model for the four scenarios and different time periods. **a**, – **c**, scenario 1 (no immigration: *v* = 0, no niche differences: *β*_*fi*_ = *β*_*ff*_), **d**, – **f**, scenario 2: adds niche differences (*v* = 0, *β*_*fi*_ < *β*_*ff*_), **g** – **i**: scenario 3 adds immigration (*v* = 0.1, *β*_*fi*_ = *β*_*ff*_), and **g**, – **i**, scenario 4 adds niche differences and immigration (*v* =0.1, *β*_*fi*_ < *β*_*ff*_). Results are shown for *t* = 200 (1000 years, left), *t* = 1000 (= 5000 years; middle), and *t* = 2000 (*t* = 10,000 years; right). The red line is the one-to-one line.

**Extended Data Figure 6.**
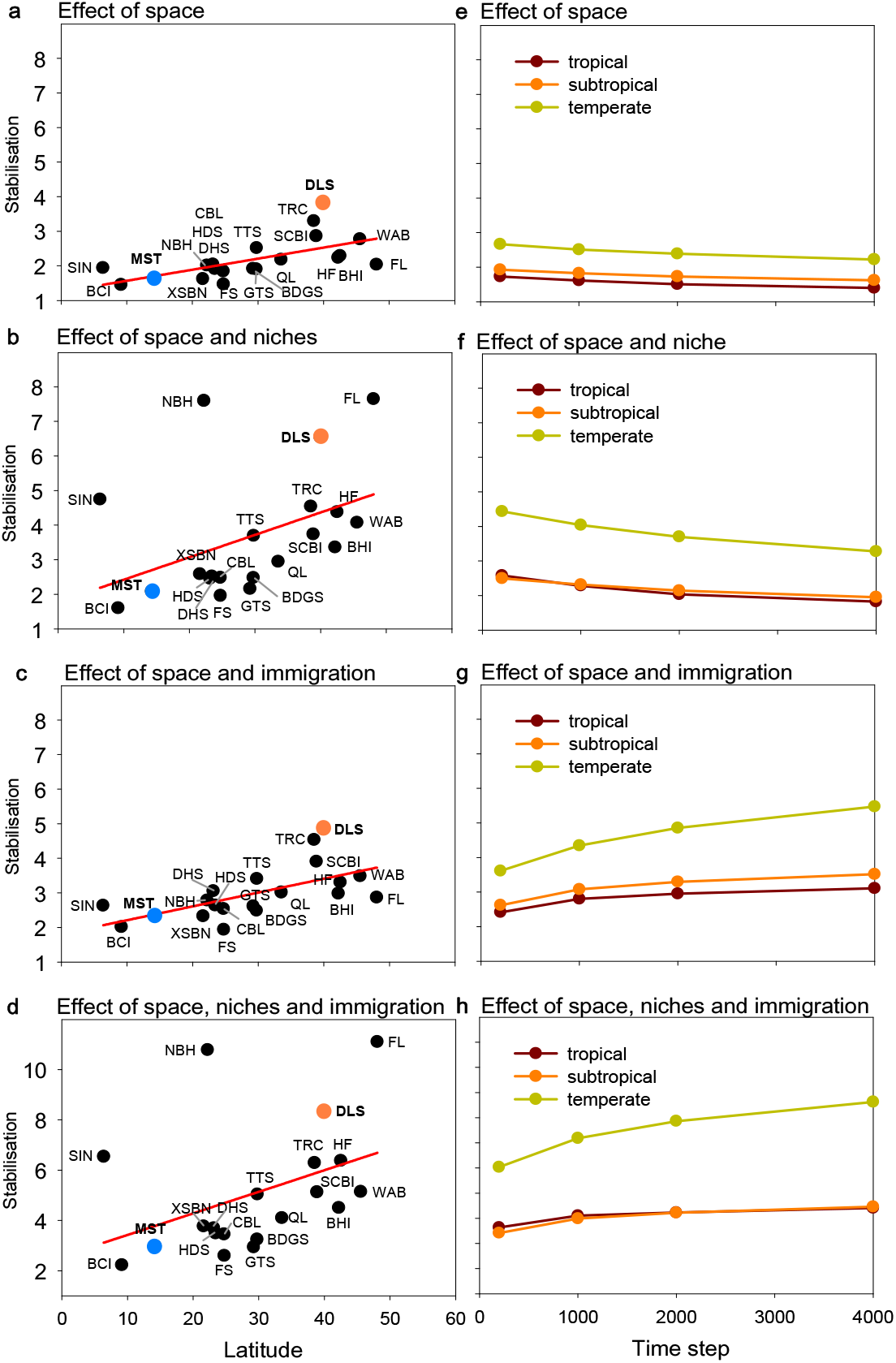
Stabilization (the factor by which the extinction risk is reduced) by different non-neutral mechanisms through time. Latitudinal variation in the reduction 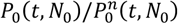 of the extinction risk, as influenced by spatial structure, niche differences and immigration. **a**, mean stabilisation at year 1000, per forest plot for scenario 1 (no immigration: *v* = 0, no niche differences: *β*_*fi*_ = *β*_*ff*_) that represents the effect of spatial structure, **b**, same as a, but for scenario 2 that adds niche differences (*v* = 0, *β*_*fi*_ < *β*_*ff*_), **c**, same as a, but for scenario 3 adds immigration to scenario 1 (*v* =0.1, *β*_*fi*_ = *β*_*ff*_), **d**, scenario 4 adds niche differences and immigration (*v* =0.1, *β*_*fi*_ < *β*_*ff*_). **e** – **h**: Changes in stabilisation through time, separately for tropical, sub-tropical and temperate forests. **e**, scenario 1, **f**, scenario 2, **g**, scenario 3, and **h**, scenario 4. The initial abundance was *N*_o_ = 50 individuals. In scenarios with niche differences we assumed that more closely related species competed more strongly. The birth-death models were parameterized for 720 species of the 21 ForestGEO plots. We excluded the temperate forest at CBS because the exponent *b*_*f*_ < −1 (Extended Data Fig. 1C) produced an extinction risk higher than the neutral model. To outline the overall tendency in the data we fitted in panels a) to d) a linear regression to the data.

**Extended Data Figure 7.**
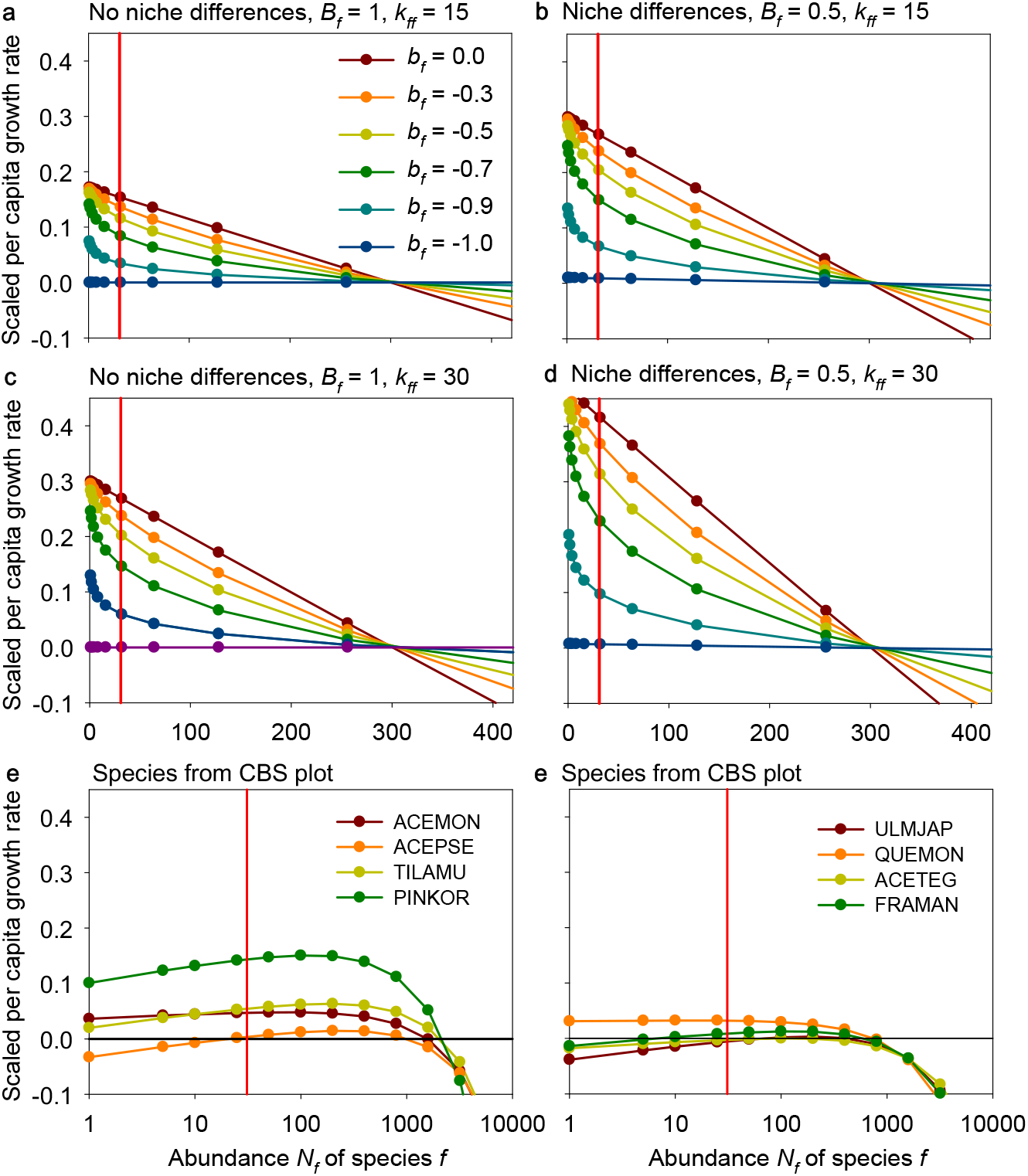
Dependency of the per capita population growth rate on the exponent of the aggregation-abundance relationship. **a**, Examples for the influence of the exponent *b*_*f*_ on the per capita population growth rate for typical parameters 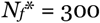, *N*_*s*_ = 31, *J* = 20,000, *r*_*f*_ = 0.1, *k*_*ff*_ = 15, *k*_*fh*_ = 0.96, *B*_*f*_ = 1 and *c* = 0.000189. **b**, same as a, but for niche differences (*B*_*f*_ = 0.5). **c**, and **d**, same as a) and b), but larger aggregation (i.e., *k*_*ff*_ = 30). **e**, and **f**, examples for the CBS plot for scenario 2 with niche differences (i.e., *B*_*f*_ < 1). We assumed that individuals compete at the individual scale more strongly if they phylogenetically more similarity. The CBS plot shows a power law exponent of *b*_*f*_ = −1.077, which leads to unstable dynamics if con- and heterospecifics compete equally at the individual scale. However, niche differences allow a few species (*Acer mono* (ACEMON), *Tilia amurensis* (TILAMU), *Pinus koraiensis* (PINKOR), *Ulmus japonica* (ULMJAP), *Quercus mongolica* (QUEMON)) to fulfil the deterministic invasion criterion (equation 11a). They are all abundant species that show weak aggregation. The vertical red lines indicates the value of the average individual fitness (at *N*_*s*_ = 31) that determines the extinction risk of a population with initially 50 individuals after 1000 years.

**Extended Data Figure 8.**
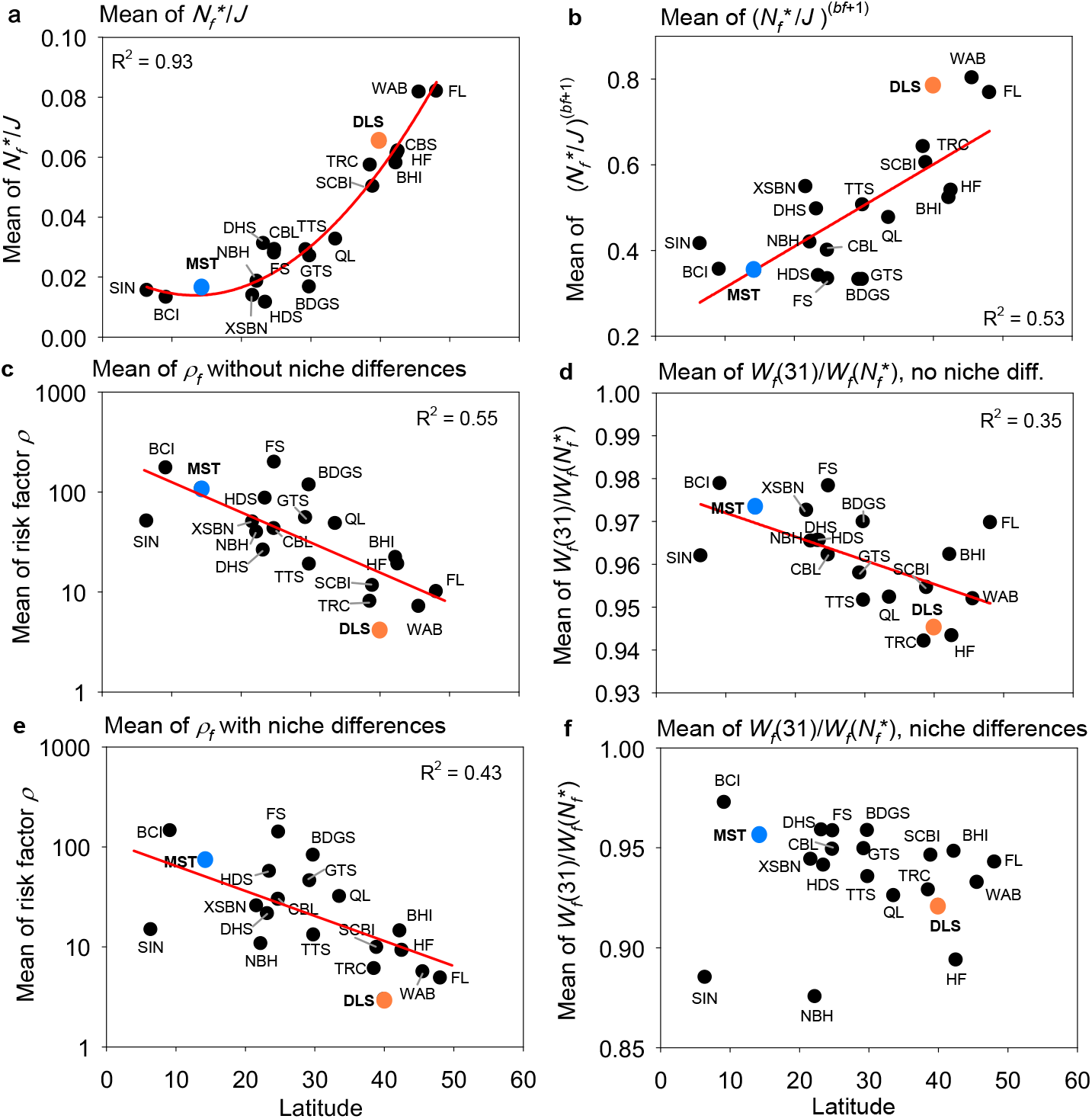
Which properties of a species lead to a low extinction risk? The panels show different quantities that drive the average population growth 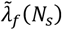 at the best fitting typical abundance *N*_*s*_, which determines the extinction risk (equation 17). **a**, the mean relative abundance 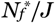, which drives the risk factor *ρ*_*f*_, averaged over all species of a given plot shown in dependence on latitude, **b**, same as a), but 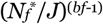 as it appears in equation 16a, **c**, the mean of the risk factor *ρ*_*f*_ taken over all focal species in each plot, in dependence on latitude, **d**, the mean of the ratio 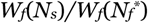 for *N*_*s*_ = 31 (eq. 16a) that drives the average individual fitness (eqs. 9a, 17) and thereby the extinction risk (equation 2), **e**, same as c), but for niche differences, **f**, same as d), but for niche differences. To outline the overall tendency in the data we fitted in panels a) a polynomial regression of order 2 and in panels b) and d) a linear regression, and in panels c) and e) an exponential regression.

**Extended Data Figure 9.**
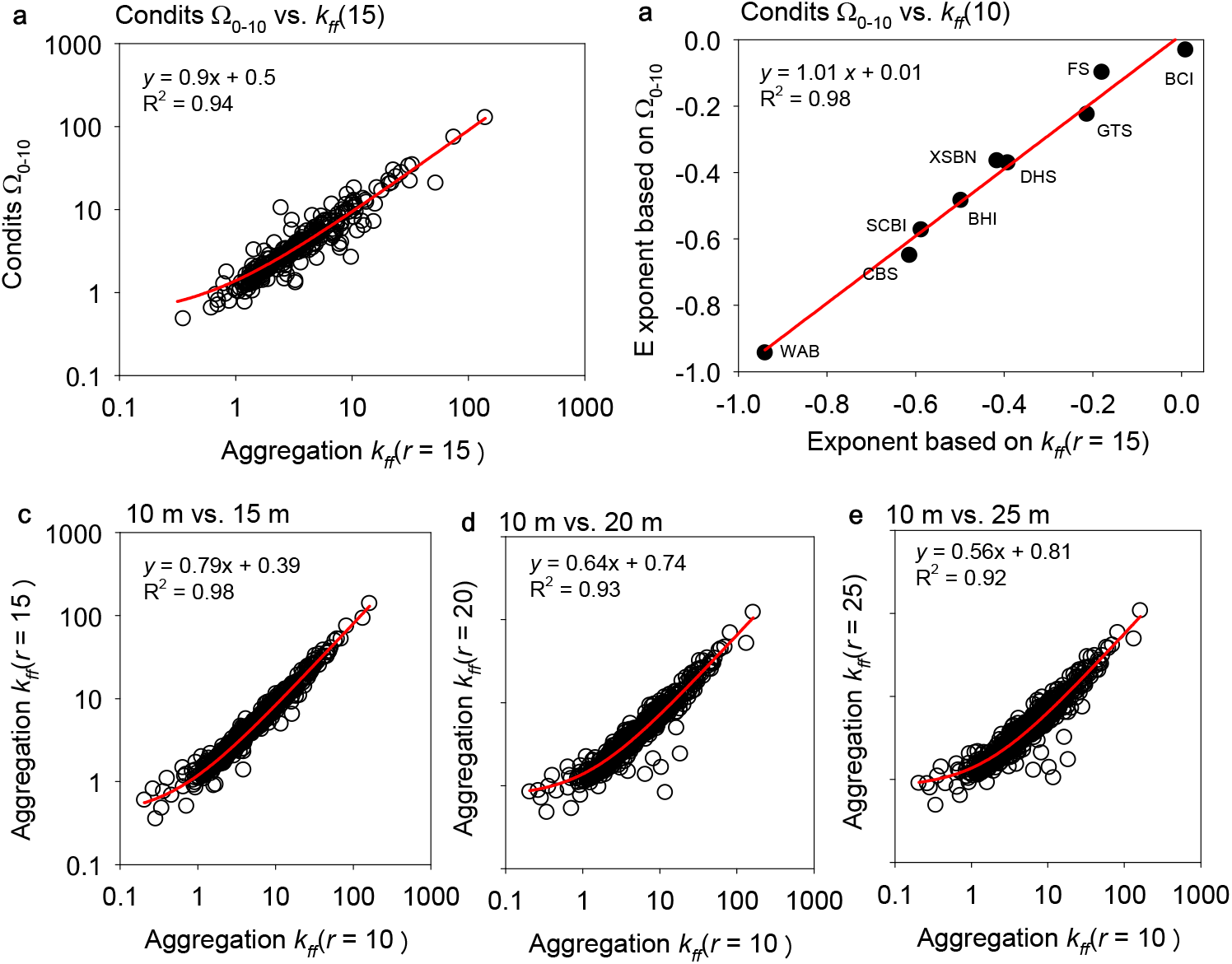
Sensitivity of the aggregation-abundance relationships to the measure of aggregation. **a**, Condits’s Ω_1-10_ aggregation measure for the species analysed in Wiegand et al. (2021) plotted over the corresponding values of the distance-weighted aggregation measure *k*_*ff*_(*r* = 15) used in the present study, **b**, exponents of the aggregation-abundance power-law derived with Condits’s Ω_1-10_ plotte over the corresponding values of the distance-weighted aggregation measure *k*_*ff*_(*r* = 15), **c**, comparison of the distance-weighted aggregation measure *k*_*ff*_(*r*) for different neighborhood radii 10m vs. 15m, **d**, comparison of the distance-weighted aggregation measure *k*_*ff*_(*r*) for different neighborhood radii 10m vs. 20m, **e**, comparison of the distance-weighted aggregation measure *k*_*ff*_(*r*) for different neighborhood radii 10m vs. 25m. To outline the overall tendency in the data we fitted linear regressions to the data.

**Extended Data Figure 10.**
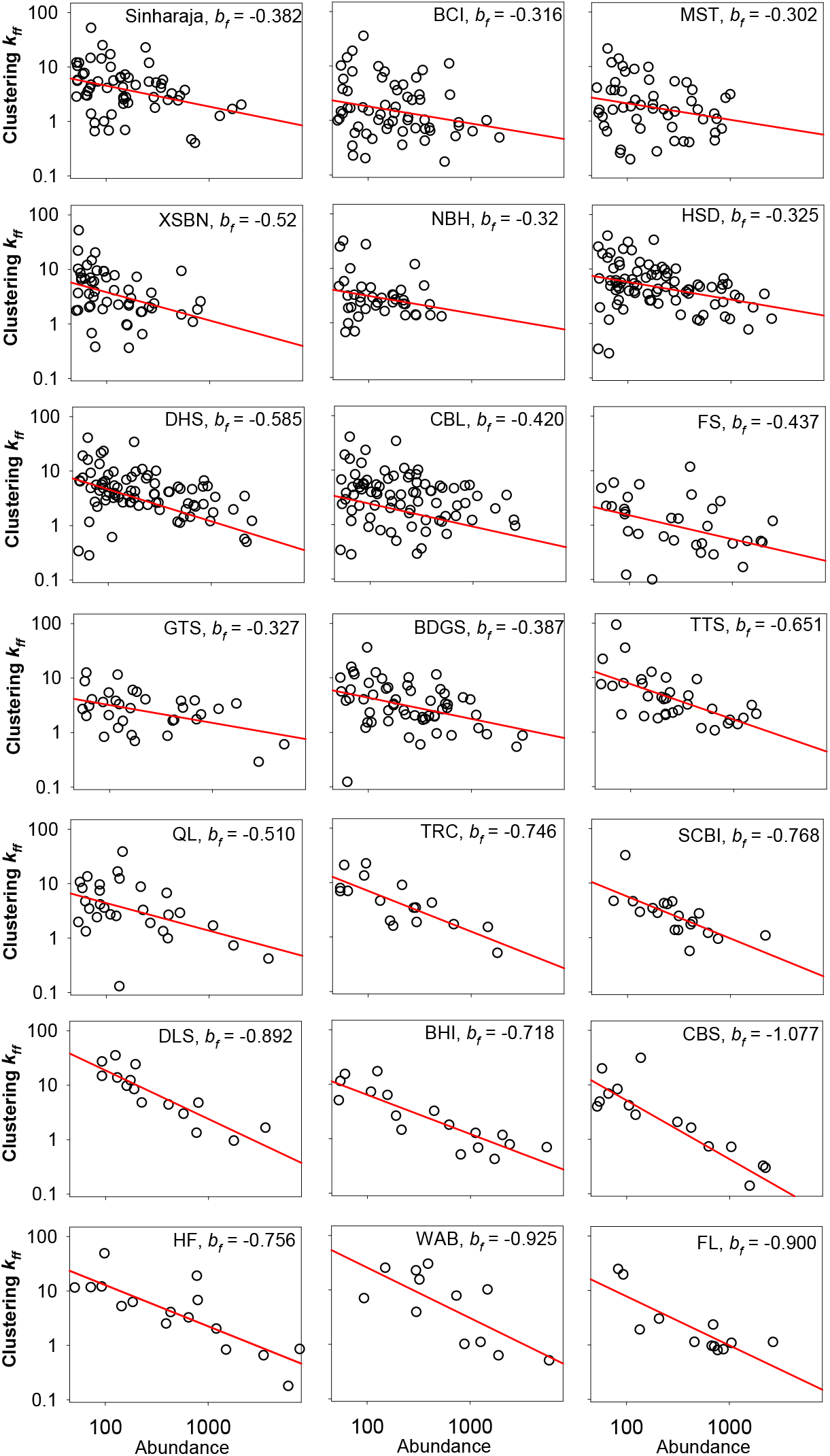
Aggregation − abundance relationships for the 21 Forest-GEO plots. The panels show the relationships between corrected aggregation *L*(*N*_*f*_)=(*k*_*ff*_ – *k*_*fh*_) (y-axis) and abundance *N*_*f*_ (x-axis) (equation 8). The value of *b*_*f*_ is the slope of the power law *L*(*N*_*f*_) = *a*_*f*_ 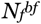 (red line), estimated by linear regression analysis of log(*k*_*ff*_ – *k*_*fh*_) over log(*N*_*f*_). We used for the analysis focal species *f* with more than 50 individual. For plot acronyms see Supplementary Table 1.

**Supplementary Table 1:** Characteristics of the selected forest plots.

| Plot | Acronym | Latitude | Elevation | Forest type<sup>†</sup> | No. species | Individuals $\geq 10\text{cm dbh}$ | Species $\geq 10\text{cm dbh}$ | Focal species<sup>#</sup> | Area (ha) | Mortality rate<sup>††</sup> | Prop. zoochory<sup>§</sup> | Prop. AM<sup>§</sup> | Prop. AM + zoochory<sup>§§</sup> |
| --- | --- | --- | --- | --- | --- | --- | --- | --- | --- | --- | --- | --- | --- |
| Sinharaja | SHJ | 6.40 | 500 | trop. | 239 | 17070 | 167 | 59 | 25 | 0.087 | 0.75 | 0.79 | 0.71 |
| BCI | BCI | 9.15 | 140 | trop. | 304 | 20647 | 238 | 63 (-10) | 50 | 0.113 | 0.89 | 1.00 | 0.91 |
| MoSingto | MST | 14.26 | 770 | trop. | 262 | 15665 | 214 | 52 (-2) | 30 | 0.136 | 0.88 | 0.91 | 0.80 |
| Xishuangbanna* | XSBN | 21.61 | 789 | trop. | 468 | 12263 | 339 | 52 | 20 | 0.14 | 0.81 | 0.85 | 0.67 |
| Nabanhe | NBH | 22.23 | 932 | trop. | 290 | 8512 | 221 | 40 (-2) | 20 | - | 0.78 | 0.86 | 0.67 |
| Heishiding | HSD | 23.45 | 567 | trop-subtr. | 245 | 30992 | 148 | 82 | 50 | 0.1 | 0.84 | 0.81 | 0.68 |
|  |  |  |  |  |  |  |  |  |  | - |  |  |  |
| Dinghushan* | DHS | 23.16 | 350 | subtr. | 210 | 11906 | 131 | 30 | 20 | 0.18 | 0.87 | 0.80 | 0.77 |
| Chebaling | CBL | 24.72 | 495 | subtr. | 228 | 10159 | 148 | 32 | 20 | - | 0.78 | 0.84 | 0.65 |
| Fushan* | FS | 24.76 | 667 | subtr. | 110 | 19261 | 77 | 33 (-1) | 25 | 0.071 | 0.85 | 0.81 | 0.68 |
| Gutianshan* | GTS | 29.25 | 581 | subtr. | 159 | 18215 | 109 | 33 | 24 | 0.09 | 0.73 | 0.68 | 0.48 |
| Badagongshan | BDGS | 29.77 | 1406 | subtr. | 232 | 25121 | 186 | 57 | 25 | 0.06 | 0.67 | 0.77 | 0.54 |
| Tiantong | TTS | 29.81 | 454 | subtr. | 154 | 14967 | 107 | 34 | 20 | 0.08 | 0.68 | 0.79 | 0.55 |
| Qinling* | QL | 33.54 | 1431 | temp.-subtr. | 121 | 11698 | 84 | 29 | 25 | - | 0.45 | 0.61 | 0.29 |
| Tyson Research Center | TRC | 38.52 | 203 | temp. | 46 | 6520 | 38 | 17 | 20 | 0.085 | 0.82 | 0.47 | 0.29 |
| SCBI | SCBI | 38.89 | 330 | temp. | 65 | 8165 | 49 | 19 | 25 | 0.08 | 0.68 | 0.42 | 0.17 |
| Donglingshan | DLS | 39.96 | 1400 | temp. | 51 | 9558 | 39 | 15 | 20 | 0.04 | 0.20 | 0.60 | 0.13 |
| Baihua* | BHI | 42.22 | 793 | temp. | 63 | 17197 | 34 | 17 | 24 | 0.07 | 0.24 | 0.53 | 0.12 |
| Changbaishan* | CBS | 42.38 | 801 | temp. | 52 | 9995 | 34 | 16 | 25 | 0.05 | 0.25 | 0.69 | 0.13 |
| Harvard Forest | HF | 42.54 | 354 | temp. | 55 | 23901 | 34 | 16 | 35 | 0.14 | 0.50 | 0.25 | 0.13 |
| Wabikon* | WAB | 45.55 | 501 | temp. | 36 | 14021 | 22 | 12 (-2) | 25.2 | 0.04 | 0.00 | 0.42 | 0.00 |
| Fenglin | FL | 48.08 | 447 | temp. | 46 | 8601 | 23 | 12 | 30 | 0.11 | 0.08 | 0.50 | 0.00 |

\* Bar code phylogeny, genus level phylogenies were used for all other plots; <sup>†</sup> trop.: tropical forests, subtr.: subtropical forests, temp.: temperate forests; <sup>#</sup> The numbers give the number of focal species with at least 50 individuals with dbh $\geq 10\text{cm}$ , we excluded species with $k_{ff} < k_{fh}$ (the numbers in parenthesis) and two species at the Wabikon forest located in a patch of successional forest that was logged approximately 40 yr ago; <sup>††</sup> The average mortality rate for trees with dbh $> 10\text{ cm}$ in the plot. Assuming approximate equilibrium, we used the values of the mortality rate to parameterize the unknown per capita recruitment rates $r_f$ . We used a recruitment rate of 0.1 for the 3 plots without mortality data. The mortality rate of the TRC plot was estimated from a 13.4 ha section of the 20 ha plot; <sup>§</sup> Proportion of species showing mostly animal seed dispersal; <sup>§§</sup> Proportion of species with arbuscular mycorrhizal association.

**Supplementary Table 2:**
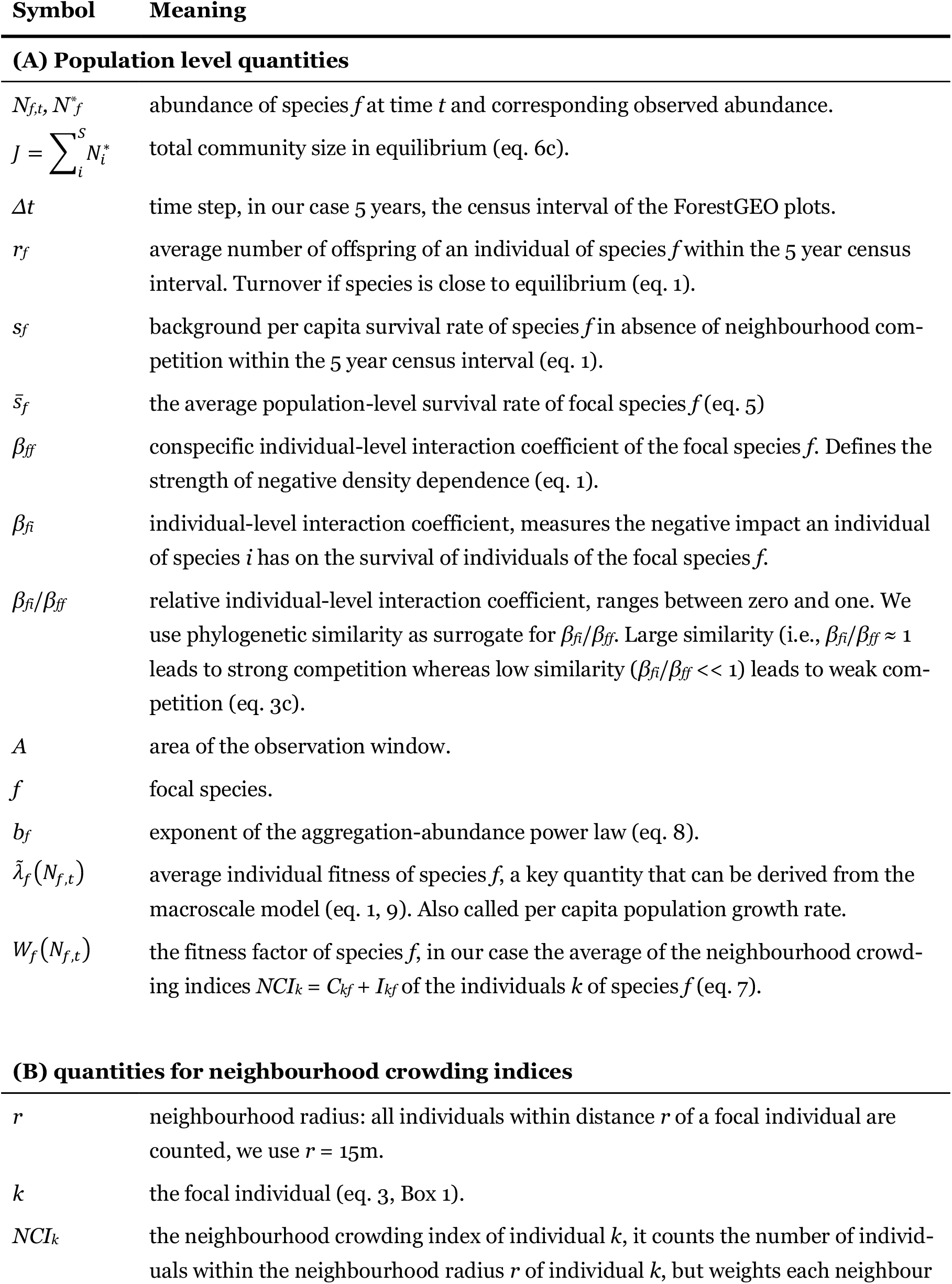
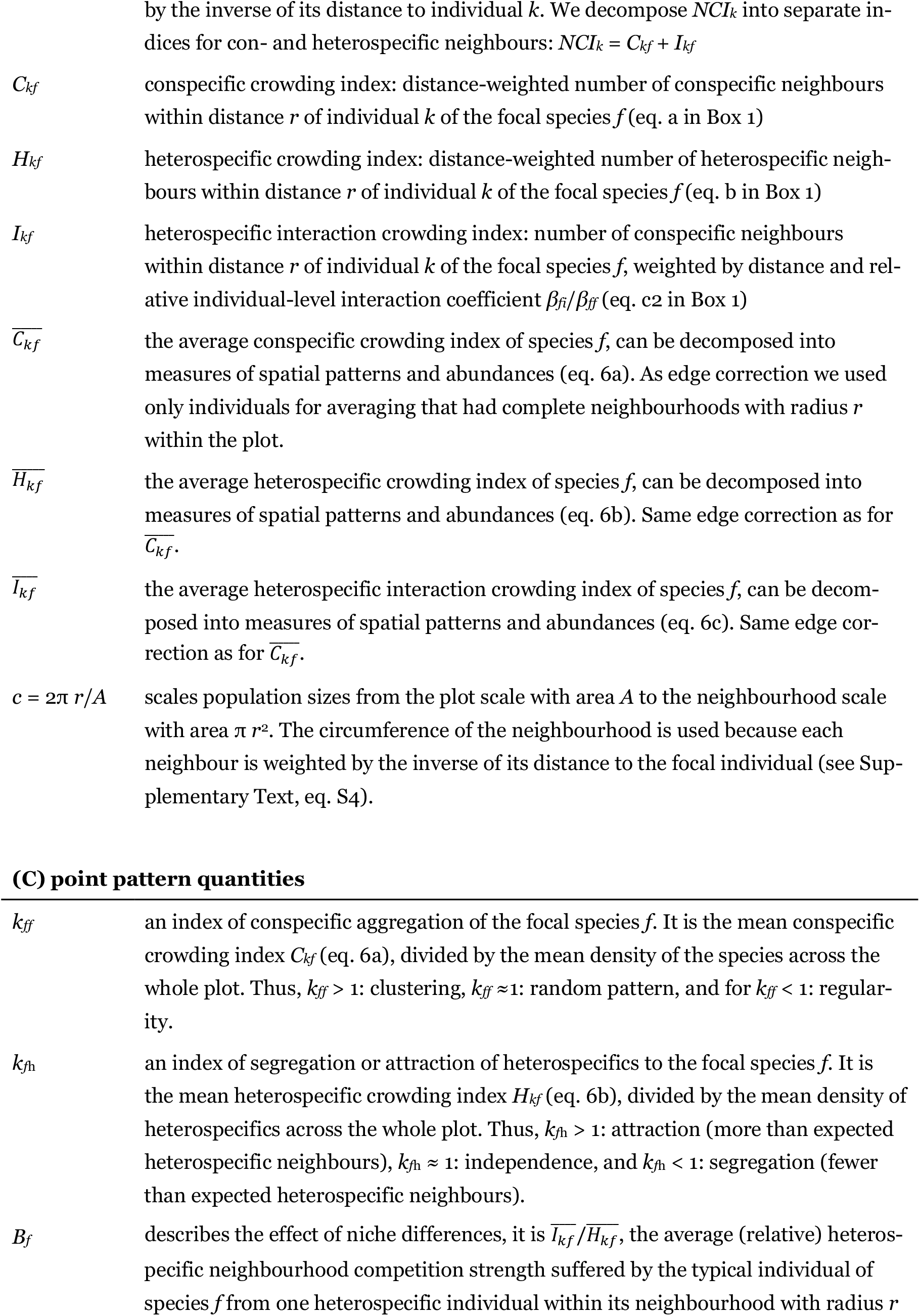
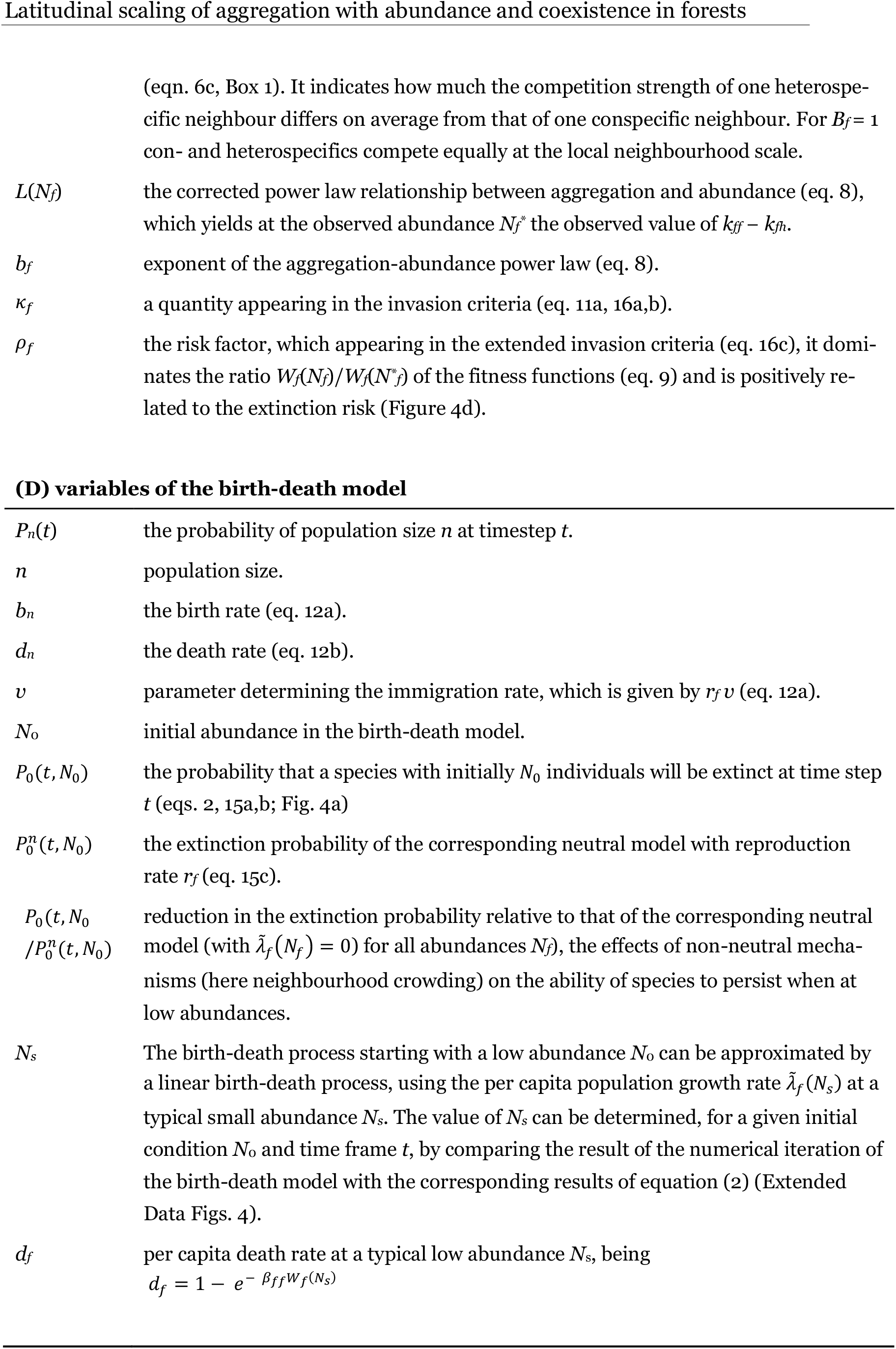
Explanation of important mathematical symbols.

## Supplementary Text

### Studies investigating the aggregation-abundance relationship

We list here studies that related aggregation in the spirit of Condit et al. (2000) to abundance, they generally found that Condit’s Ω increased with decreasing abundance. Unfortunately, most of these studies showed graphs that correlated Ω or SES(Ω) (but not ln Ω) with ln(abundance), but often for several size classes together.

## Details of the spatial analysis

### Decomposition of average crowding indices

The task is to relate the neighbourhood-scale averages of the crowding indices 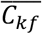 and 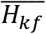 shown in Box 1 to the population-scale abundances *N*_*f*_ and measures of spatial patters (i.e., conspecific aggregation and heterospecific segregation). We solve this up-scaling problem using point-process theory (Wiegand and Moloney 2014; Illian et al. 2008). Before we explain how we obtain equations 6a and 6b in the main text, we present some basics of spatial point process theory and a simpler example of a crowding index (Wiegand et al. 2021) that leads to the aggregation measure Ω_0–10_ presented in Condit et al. (2000).

In the simplest case, an index of conspecific crowding simply counts the number of neighbours within distance *r* of the focal individual (Wiegand et al. 2021). This leads to Ripley’s *K*-function, a well-known quantity in spatial statistics. The *K*-function can be estimated as the expected number of neighbours occurring within distance *r* of the typical individual, divided by *λ*_*f*_, the overall density of a species in the plot (i.e., *λ*_*f*_ = *N*_*f*_ /*A*, the number *N*_*f*_ of individuals divided by the area *A* of the plot). Thus, in the simplest case, the conspecific crowding index is given by *λ*_*f*_ *K*(*r*).

Spatial aggregation describes the extent to which trees of the same species tend to occur in spatial clusters and is usually defined as the average density of conspecific trees in circular neighbourhoods around each individual tree, divided by the mean density of the species across the whole plot (Condit et al. 2000; Wiegand and Moloney 2014).Using the *K*-function, we obtain with this definition the measure of aggregation *k*_*ff*_ (*r*) = *K*(*r*)/π*r*^2^. If there is an excess of neighbours within distance *r* (i.e., *k*_*ff*_(*r*) > 1) we have aggregation, and if there are fewer than expected neighbours, we have regularity (i.e., *k*_*ff*_(*r*) < 1).

We can now decompose our simple crowding index *λ*_*f*_ *K*(*r*) into the measure *k*_*ff*_(*r*) of aggregation and species abundance *N*_*f*_, given that *λ*_*f*_ = *N*_*f*_/*A* we obtain

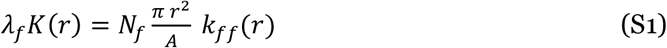

where the quantity π*r*^2^/*A* is a scaling constant that scales the number of individuals from the entire plot with area *A* to the neighbourhood with area π*r*^2^.

To decompose our crowding indices that additionally weight each individual by 1 over distance to the focal individual, we first recognize that the *K*-function is the cumulative version of the pair correlation function *g*(*r*):

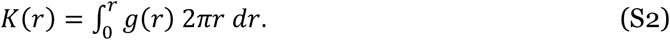

The quantity *λ*_*f*_ *g*(*r*) is the density of points within a ring with radius *r* and width *dr*, centred at the typical individual, and multiplying with the area of that ring (=2*πr dr*) yields the expected number of neighbours in this ring. Summing up over all rings up to radius *r* gives the expected number of neighbours occurring within distance *r* of the typical individual (i.e., *λ*_*f*_*K*(*r*)). The pair correlation function is normalized in a way that it has a value of *g*(*r*) = 1 if the neighbourhood density is identical to the overall density *λ*_*f*_ of individuals in the plot (= *N*_*f*_/*A*). Thus, the value of one of the pair correlation function serves as dividing line between cases of aggregation where *g*(*r*) > 1 (i.e., the typical individual has more neighbours than expected at distance *r* by a random distribution) and cases of regularity where *g*(*r*) < 1 (i.e., the typical individual has less neighbours than expected at distance *r* by a random distribution).

Taking advantage of equation S2, we write the distance-weighted conspecific crowding index as

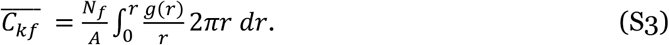

We can do this since the pair correlation function looks only at neighbours at distance *r* of the typical individual. The *r* cancels, and with small rearrangements and adding factor *r* (1/*r*) we obtain the decomposition of the conspecific crowding index (equation 6a in the main text):

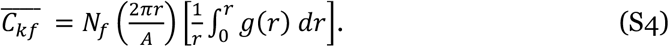

into abundance *N*_*f*_, a scaling constant *c* = 2π*r*/*A*, and our measure of aggregation

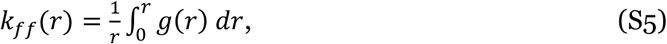

which is the average of the pair correlation function up to distance *r*. The measure *k*_*ff*_(*r*) has the desired properties of a measure of aggregation, if the average neighbourhood density of conspecific trees equals the mean density of the species across the whole plot, we obtain *g*(*r*) = 1 and therefore *k*_*ff*_(*r*) =1, if the number of neighbours is larger we obtain *g*(*r*) > 1 (= aggregation) and if the number of neighbours is smaller we obtain *g*(*r*) < 1 (= regularity).

In the same way we obtain the scaling relationship for the heterospecific crowding index 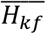 that counts the (distance-weighted) number of heterospecific neighbours within distance *r* of the typical individual of the focal species *f*. Here we use for the decomposition the bivariate pair correlation function *g*_*fh*_(*r*), the density of heterospecific neighbours at a small distance interval (*r*−*dr*/2, *r*+*dr*/2) of the typical individual of the focal species *f*, divided by the overall density *λ*_*h*_ of heterospecifics:

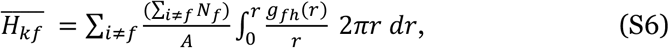

which results in the decomposition (equation 6b):

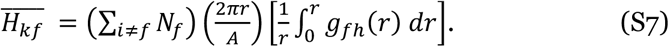

into the sum of the abundances of heterospecifics ∑_*i*≠*f*_ *N*_*f*_, the scaling constant *c* = 2π*r*/*A*, and our measure of segregation

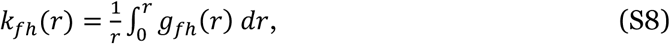

the average of the bivariate pair correlation function up to distance *r*. The measure *k*_*fh*_(*r*) has the desired properties of a measure of segregation, defined as the degree of reduction of heterospecific neighbours, compared to the expectation of a independent species distribution: if the focal species is located independently on heterospecifics, we obtain *g*_*fh*_(*r*) = 1 and therefore *k*_*fh*_(*r*) =1, if the number of heterospecific neighbours is smaller than expected we obtain *k*_*fh*_(*r*) < 1 (= segregation) and if the number of heterospecific neighbours is larger we obtain *k*_*fh*_(*r*) > 1 (= attraction).

To obtain the scaling relationship for the crowding index 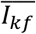 that weight additionally to 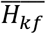 each heterospecific neighbour by its relative interaction strength *β*_*fi*_/*β*_*ff*_ we first define the quantity

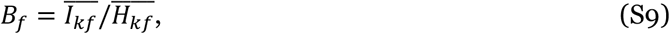

the average competition strength of one heterospecific neighbour relative to that of one conspecific. We found that the quantity *B*_*f*_ is for species rich forests at or near a stationary state in good approximation a species-specific constant (see Supplementary Text in Wiegand et al. 2021). Thus, we can apply a mean-field approximation (O’Dwyer and Chisholm 2014; Fung et al. 2022) where the species-specific competition strengths of heterospecifics can be replaced by an average heterospecific competition strength, the quantity *B*_*f*_. The *B*_*f*_ summarizes the emerging effects of the individual-level interaction coefficients *β*_*fi*_/*β*_*ff*_ at the population-level. Note that this approximation does not mean that we ignore differences in pairwise interaction strength at the individual level, where competitive interactions occur, but the approximation tells us that the effect of existing differences in pairwise interactions strengths between species average out at the community level.

Using equation S9, we can decompose the crowding index 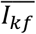 together with equation S7 into

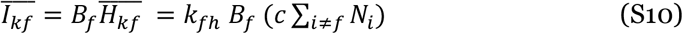

Given that strong local density dependence causes approximate zero-sum dynamics, where the total number *J* of individuals remains constant, we can approximate the number of heterospecifics in equation S10 by *J* − *N*_*f*_ and obtain the final decomposition (eq. 6c):

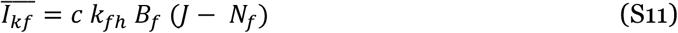

Using these approximations decouples the multispecies dynamics and allows us to investigate the dynamics of individual species in good approximation.

When estimating the crowding indices and the indices of spatial patterns from the data, we need to conduct edge correction. This is because individuals too close to the edge of the plot do not have complete neighbourhoods with radius *r* inside the plot. We can therefore not evaluate their number of con- and heterospecific neighbours. To avoid bias, we excluded these individuals.

## Model equations for alternative models

In equation 7 we present a simple macroscale model where neighbourhood crowding governs the survival of individual adult trees. However, the neighbourhood crowding can also affect reproduction (i.e., the number of offspring produced per adult) or establishment of offspring (i.e., whether or not an offspring placed at a given location can establish). In the following we present several alternative models with neighbourhood crowding in adult survival, offspring survival and reproduction.

### Alternative Model A1: Density dependence in reproduction due to neighbourhood crowding

The spatially-enriched macroscale model for the per capita growth rate of species *f* with neighbourhood crowding affecting reproduction is given by

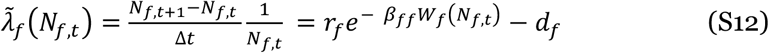

where *r*_*f*_ is the maximal reproduction rate (at a location without crowding competition were the fitness factor *W*_*f*_ = 0), *d*_*f*_ is the average per capita mortality rate of adults obtained from the data, *β*_*ff*_ the individual-level conspecific interaction coefficients of species *f*, and *W*_*f*_(*N*_*f,t*_) the fitness factor of equations (7). Note that the crowding indices count, similar to the case were neighbourhood crowding affects survival, conspecific and heterospecific neighbours around adult individuals. Assuming approximate equilibrium, we obtain from equation S12 the unknown value of the individual-level conspecific interaction coefficients 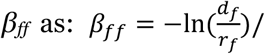 *W*_*f*_(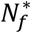).

The corresponding full birth-death model is therefore given by

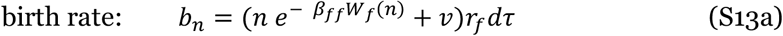

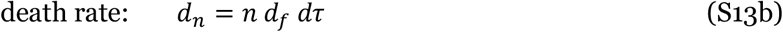

and the approximation of the birth rate at a typical small abundance *N*_s_ is given by

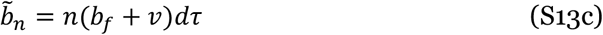

where 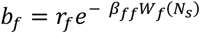. Note that 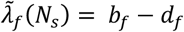. The extinction risk for the linear birth-death process (equations 13b,c) is given by

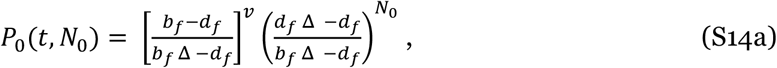

where 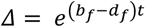. We now define 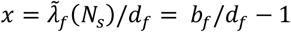 and *T* = *d*_*f*_*t* being the time scaled by the mean mortality rate, and obtain Δ = e^*xT*^ and

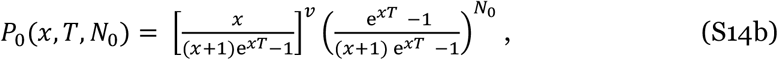

Equation S14b can be directly compared to equation 2 in the main text.

### Alternative model A2: Density dependence in establishment of offspring

The equations for alternative model A1 above apply also for the case where neighbourhood crowding affects the establishment of offspring after it is placed at a given location. This is because in model A1 the number of established recruits is reduced by distributing less recruits that then always establish (i.e., on average 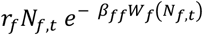 recruits are distributed), whereas in model A2 *r*_*f*_ *N*_*f,t*_ recruits are distributed on average, and then, depending on their neighbourhood, they can establish with probability 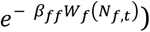. However, in model A2, the crowding indices need to count conspecific and heterospecific neighbours around the locations of the tentative offspring. For example, the conspecific crowding index *C*_*kf*_ is then given by the (distance-weighted) number of conspecific adults located within distance *r* of an offspring individual *k*. This somewhat complicates the estimation of the crowding indices and indices of spatial patterns, because only the fraction *r*_*f*_ *N*_*f*_ of focal individuals (i.e., tentatively placed recruits) can be used to estimate the average crowding and pattern indices, whereas in the alternative model A1 (and the model of equation 7) we have 10 time more focal individuals (i.e., adults) to estimate the indices, if *r*_*f*_ (or *d*_*f*_) = 0.1.

In the alternative model A2 we expect a trade-off behaviour of stabilisation in response to changing the rules of placement of offspring relative to their parents. In the one extreme, when offspring is placed close to their parents, a negative relationship between aggregation and abundance will arise as in our model (eq. 7) and reduce stabilisation. However, in the other extreme, when offspring is placed in an aggregated way away from their parents, the corresponding aggregation index (which is now not adult aggregation, but aggregation of adults around tentative offspring) will approach values of one because offspring is placed independently on the locations of their parents. This leads to low stabilisation. We expect that maximal stabilisation will be reached if a certain proportion of offspring remains close to the parents (e.g., *b*_*f*_ = −0.4) and/or the random cluster centres around which offspring is placed persist more than one census period.

### Alternative model A3: Density dependence in reproduction and mortality due to neighbourhood crowding

This spatially-enriched macroscale model corresponds to the classical Lotka-Volterra model and is given by

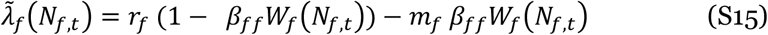

where *r*_*f*_ and *m*_*f*_ are the per capita reproduction and mortality rates without crowding competition (i.e., *W*_*f*_(*N*_*f,t*_) = 0), respectively, and *W*_*f*_(*N*_*f,t*_) is the fitness factor of equations (7). Assuming approximate equilibrium, we obtain from equation S15 the unknown value of the individual-level conspecific interaction coefficients 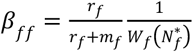.

The corresponding full birth-death model is therefore given by

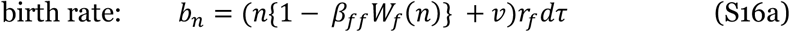

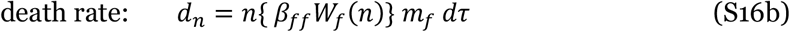

and we approximation of the birth and death rate at a typical small abundance *N*_s_ is given by

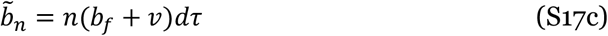

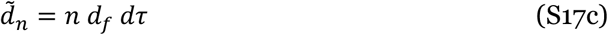

where *b*_*f*_ = *r*_*f*_{1 - *β*_*ff*_*W*_*f*_(*N*_*s*_)} and *d*_*f*_ = *m*_*f*_ {*β*_*ff*_*W*_*f*_(*N*_*s*_)}. Note that 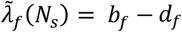. The extinction risk for the linear birth-death process (equations 13b,c) is given by

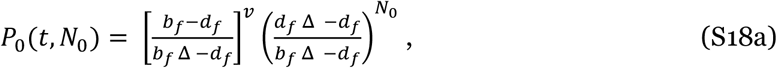

where 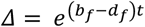. With the definitions 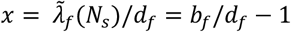 and *T* = *d*_*f*_*t* we obtain Δ = e^*xT*^ and therefore as in equation S14b:

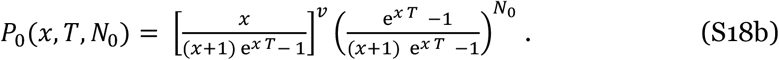

Note that in this case the scaled time *T* depends via the death rate *d*_*f*_ on the typical small abundance *N*_s_.

## Spatially-explicit simulation model

### Model description (adapted from Wiegand et al. 2021)

The individual-based simulations serve two main purposes; first, they are used to verify that the observed patterns (i.e., the abundance-aggregation power law) can emerge in principle from the minimal mechanisms included into the macroscale model, and they are used as “known data” to test if the parameterized birth-death model (equations 12 and 13) is able to predict the simulated abundance distribution correctly.

The model is an individual-based and stochastic implementation of the spatial multi-species model (equations 1, 7a), and simulates the dynamics of a community of initially *S* tree species (in our case *S* = 80) in a given plot of a homogeneous environment (in our case 200 ha) in 5-year timesteps adapted to the ForestGEO census interval. Only reproductive (adult) trees are considered, but no size differences.

During a given 5 yr timestep the model simulates first stochastic recruitment of reproductive trees and placement of recruits, and second, stochastic survival of adults (equation 4) depending on the neighbourhood crowding indices for conspecifics (*C*_*kf*_) and heterospecifics (*H*_*kf*_ or *I*_*kf*_ depending on the scenario) (but excluding recruits)(equation 3). In the next timestep the recruits count as reproductive adults and are subject to mortality. No immigration from a metacommunity is considered. To avoid edge effects torus geometry is assumed.

The survival probability of an adult *k* of species *f* is given by equation (4) and yields *s*_*kf*_ = *s*_*f*_ *exp*(– *β*_*ff*_ (*C*_*kf*_ + *I*_*kf*_)). The two neighbourhood indices *C*_*kf*_ and *I*_*kf*_ describe the competitive neighbourhood of the focal individual *k* and sum up all con-specific and heterospecific neighbours within distance *r*, respectively, but weight them by the inverse of it distance to the focal individual *k* and by the relative individual-level interaction coefficients *β*_*fi*_/*β*_*ff*_. If con- and heterospecifics compete equally at the individual-scale, we have *β*_*fi*_= *β*_*ff*_.

Each individual produces *r*_*f*_ recruits on average and their locations are determined by a type of Thomas process (Wiegand and Moloney 2014) to obtain a clustered distribution of recruits. In our model, the spatial position of the recruits is determined by two independent mechanisms. First, a proportion 1–*p*_d_ of recruits is placed stochastically around randomly selected conspecific adults (parents) by using a two-dimensional kernel function (here a Gaussian with variance *σ*^2^). This is the most common way in most spatially-explicit models to generate species clustering. Technically, we first select for each of these recruits randomly one parent among the conspecific adults and then determine the position of the recruit by sampling from the kernel. Second, the remaining proportion *p*_*d*_ of recruits is distributed in the same way around randomly placed cluster centres that are located independently of conspecific adults. We select first for each of these recruits randomly one cluster centre among the cluster centres of the corresponding species, and then determine the position of the recruit by sampling from the kernel.

### Parameterization of the simulation model

The simulation model used here is described in Wiegand et al. (2021). In contrast to Wiegand et al. (2021), we use here distance-weighting for the estimation of the crowding indices. Thus, in the source code (published as Supplementary information in Wiegand et al. 2021) we use now DistanceWeighting = 1 instead of DistanceWeighting = 0.

The simulations of the simple individual-based forest model were conducted over 25,000 yrs (= 5000 census periods) in an area of *A* = 200 ha, and comprised approximately 86,000 trees with initially 80 species. There was no immigration. The model parameters were the same for all species, and all species followed exactly the same model rules. We selected *β*_*fi*_ = *β*_*ff*_ to obtain no differences in con- and heterospecific interactions and *s*_*f*_ = 1 (no background mortality), and we adjusted the parameters *β*_*ff*_ = 0.02 and *r*_*f*_ = 0.1 to yield tree densities (430/ha) and an overall 5 year mortality rate (10%) similar to that of trees with dbh ≥ 10cm of the BCI plot^71^. The radius of the neighbourhood used to estimate the crowding indices was *r* = 15m.

To test if the birth-death model is able to correctly predict the species abundance distribution of the individual-based implementation of our model, we selected three different parameterizations of the placement of recruits that produce typical average individual fitness functions (Fig. S1a). To this end we used for the three parameterizations respective values of *σ* = 67, 15, 10m for the parameter *σ* of the Gaussian kernel that places recruits around conspecific adults or around randomly placed cluster centres. We obtain weak spatial aggregation for *σ* = 67 and stronger smaller-scale aggregation for the values *σ* =15, 10 m to mimic more realistic seed dispersal distances (Wiegand et al. 2017). The probabilities *p*_*d*_ that a new recruits was located close to one of the randomly placed cluster centres were *p*_*d*_ = 0.0125, 0.6, and 1, respectively, while the probability to be placed close to a conspecific adult was 1 - *p*_*d*_. We used in our simulations 16 random cluster centres that changed every timestep their location.

### Characteristics of the model scenarios

The first scenario represents a case of weak aggregation (σ = 67m). It results in the Latitudinal scaling of aggregation with abundance and coexistence in forests power law 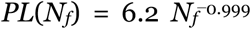 with an average individual fitness functions of almost zero for all abundances (Fig. S1a, top), and the birth-death mode predicts after 5000 census periods *Δt* a very wide probability distribution *P*_*n*_(*t*) (Fig. S1b, top). The second scenario represents realistic aggregation (σ = 15m) with recruits placed in similar proportions around their parents and randomly selected cluster centres. It results in the power law 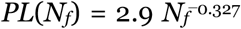 with a moderate average individual fitness functions at lower abundances (Fig. S1a, middle), and a much narrower distribution *P*_*n*_(*t*) (Fig. S1a, middle). The third scenario represents strong aggregation (σ = 10m) with recruits placed only around the randomly selected cluster centres. It results in the power law 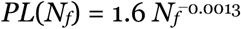 with a high average individual fitness functions at low abundances (Fig. S1a, bottom) and a narrow distribution *P*_*n*_(*t*) (Fig. S1b, bottom). In all cases, the probability distribution *P*_*n*_(*t*) predicted by the birth-death model matched the abundance distribution of the individual-based simulations after 5000 census periods well (Figure 1c).

---

**Figure S1.**
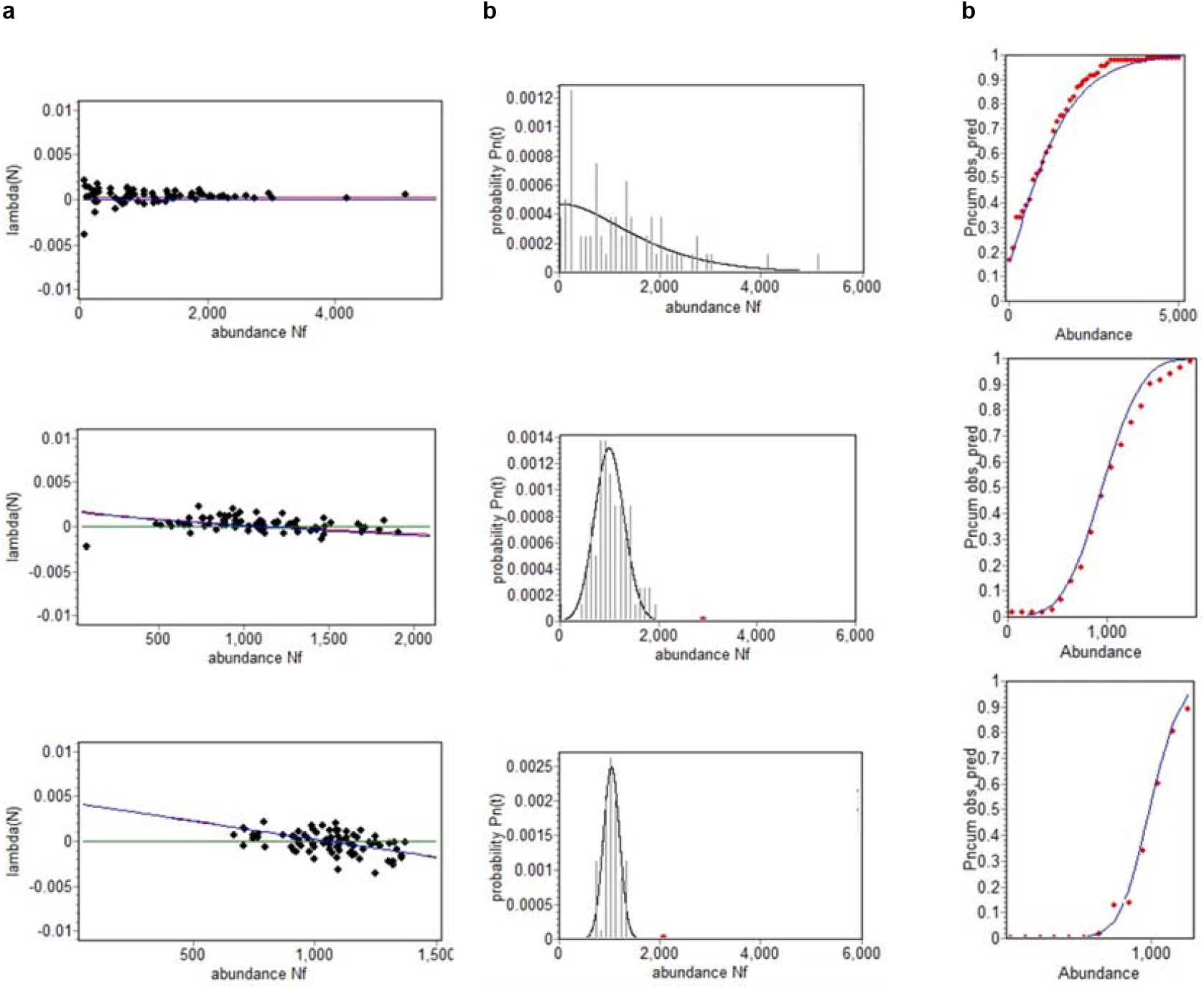
The birth-death model predicts the species abundance distribution of individual-based simulations. **a**, The average individual fitness (red-blue line) determined from the fitted power-law relationship between (*k*_*ff*_ – *k*_*fh*_) and *N*_*f*_ (Extended Data Fig. 1), the mean value of *k*_*fh*_, the observed total community size *J*, and the model parameters *β*_*ff*_ = 0.02, *r*_*f*_ = 0.1, and *s*_*f*_ = 1.0 (see methods). The black dots show the values of average individual fitness for the different species, given their observed abundance 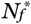, aggregation *k*_*ff*_ and segregation *k*_*fh*_, **b**, The observed species abundance distribution of the individual-based model (IMB) after 25,000 years (bars) and the corresponding prediction of the birth-death model (black line), the (scaled) probability *P*_*n*_(*t*) that the species has at time *t n* individuals, **c**, The cumulative probability distributions (red dots: simulated by IBM, blue line: predicted by birth-death model). The individual-based simulation model is a spatially explicit and stochastic implementation of the spatial multi-species model (equation 1), and simulates the dynamics of a community of initially 80 tree species in a given 200 ha plot of a homogeneous environment in 5-year time steps.

